# Probabilistic discrimination of relative stimulus features in mice

**DOI:** 10.1101/2020.12.20.423700

**Authors:** Dmitry R Lyamzin, Ryo Aoki, Mohammad Abdolrahmani, Andrea Benucci

## Abstract

Understanding how the brain computes choice from sensory information is a central question in perceptual decision-making research. From a behavioral perspective, paradigms suitable to study perceptual decision-making condition choice on invariant properties of the stimuli, thus decoupling stimulus-specific information from decision-related variables. From a neural perspective, powerful tools for the dissection of brain circuits are needed, which suggests the mouse as a suitable animal model. However, whether and how mice can perform an invariant visual discrimination task has not yet been fully established. Here, we show that mice can solve a complex orientation discrimination task where the choices are decoupled from the orientation of individual stimuli. Moreover, we demonstrate a discrimination acuity of at least 6°, challenging the common belief that mice are poor visual discriminators. We reached these conclusions by introducing a novel probabilistic choice model that explained behavioral strategies in (n = 40) mice and identified unreported dimensions of variation associated with the circularity of the stimulus space. Furthermore, the model showed a dependence of history biases on task engagement, demonstrating behavioral sensitivity to the availability of cognitive resources. In conclusion, our results reveal that mice are capable of decoupling decision-relevant information from stimulus-specific information, thus demonstrating they are a useful animal model for studying neural representation of abstract learned categories in perceptual decision-making research.

## Introduction

Most behaviorally relevant information in visual scenes is provided by the objects and relationships between them rather than the low-level visual features. Relative properties of objects, such as spatial arrangement (Krechevsky, 1938; Lashley, 1938), shape and color similarity (Martinho and Kacelnik, 2016), relative contrast (Burgess et al., 2017), and relative density or numerosity (Dakin et al., 2011), can condition behavior, which necessarily relies on the corresponding abstract, stimulus-invariant neural representations.

In perceptual decision-making research, tasks that enable neural-to-behavioral coupling need to fulfill specific requirements. First, they should rely on these relative or more abstract categories to separate the neural representation of the decision information from sensory representations, which are often encoded in the same neural populations (Akrami et al., 2018; Pho et al., 2018; Romo et al., 1999; Steinmetz et al., 2019). Second, sensory stimuli should be sufficiently complex to engage cortical computations (DiCarlo and Cox, 2007), but with known neural encoding characteristics to permit targeted neural recordings (Hubel and Wiesel, 1962; Huberman and Niell, 2011). Finally, a rich set of experimental tools for the dissection of the underlying neural circuits should be available in the animal model of choice (Abbott et al., 2020; Luo et al., 2018; Madisen et al., 2015).

Visual discrimination tasks in rodents do, in principle, fulfill all these requirements. However, previous studies have typically traded-off some of them. For instance, simple visual stimuli (e.g., light bars, contrast gratings) are easy to parametrize and their neural encoding is well characterized, but to which extent they engage cortical computations is a matter of debate (Wang et al., 2018, 2020). Similarly, visual objects or natural images are probably more effective at driving cortical computations, but the stimulus parametrization is challenging, and their neural substrate and encoding characteristics are still largely unexplored (DiCarlo and Cox, 2007; DiCarlo et al., 2012; Riesenhuber and Poggio, 2000). A viable alternative could be to condition choice on a relative property of stimuli that are easy to parametrize, and that have a well-defined cortical representation.

Here, we sought to establish if mice can learn to discriminate relative orientations, and if so, to identify their choice determinants. To this end, we developed a task, in which the animal indicates the more vertically oriented grating stimulus of two simultaneously presented. To quantify the behavior, we developed a novel probabilistic model of choice that captured choice variability and choice biases including the history-dependent ones. With the help of the model, we established how individual animals combine information about the two orientations, estimated their discrimination acuity, and demonstrated the dependency of history biases on the task engagement. We suggest that our task will allow the exploration of complex decision-making and visual-to-cognitive links in mice, particularly when studying the computation of decision in visual areas (DiCarlo and Cox, 2007), the link between neural and behavioral variability (Beck et al., 2012), and a role of heuristics and suboptimal choice strategies (Gardner, 2019).

## Results

### Relative orientation discrimination task

#### Task details

We trained transgenic mice (n = 40) in a two-alternative forced-choice (2AFC) orientation discrimination task using an automated setup, in which the animal voluntarily fixed its head to initiate an experimental session (**Fig. 1a,** top), as previously described (Aoki et al., 2017). In this task, two oriented Gabor patches were simultaneously shown on the left and right sides of a screen; to obtain water, the animals had to identify the oriented patch that was *more vertical* (n = 28; *more horizontal,* n = 12) and move it to the center of the screen by rotating a wheel manipulator (Aoki et al., 2017; Burgess et al., 2017). Crucially, because the target in most trials was not vertical, the animals had to compare the angular distance to the vertical (*verticality*) of the two orientations. The same physical stimulus could thus be a target or a nontarget in different trials, thereby making the task invariant relative to the orientation of individual stimuli (**Fig. 1a,** middle). The orientations of both stimuli (*θ_L_,θ_R_*) were sampled at random from angles between −90° and 90° with a minimal angular difference of 9° (3° for one animal), with positive angles corresponding to clockwise and negative to counterclockwise orientations relative to vertical (**Fig. 1a,** bottom). We used this 9° spacing for most animals to sample a high number of responses for every angle condition, which was important for subsequent imaging experiments (not shown in this study). We analyzed a total of 1,313,355 trials, ranging from 4591 to 82,065 per animal, with an average of 32,834 ± 2962 trials per animal (mean ± s.e.), in 256 ± 22.28 sessions of 128.02 ± 1.34 trials each (**Supplementary Fig. 1, Supplementary Table 1**).

**Figure 1.**
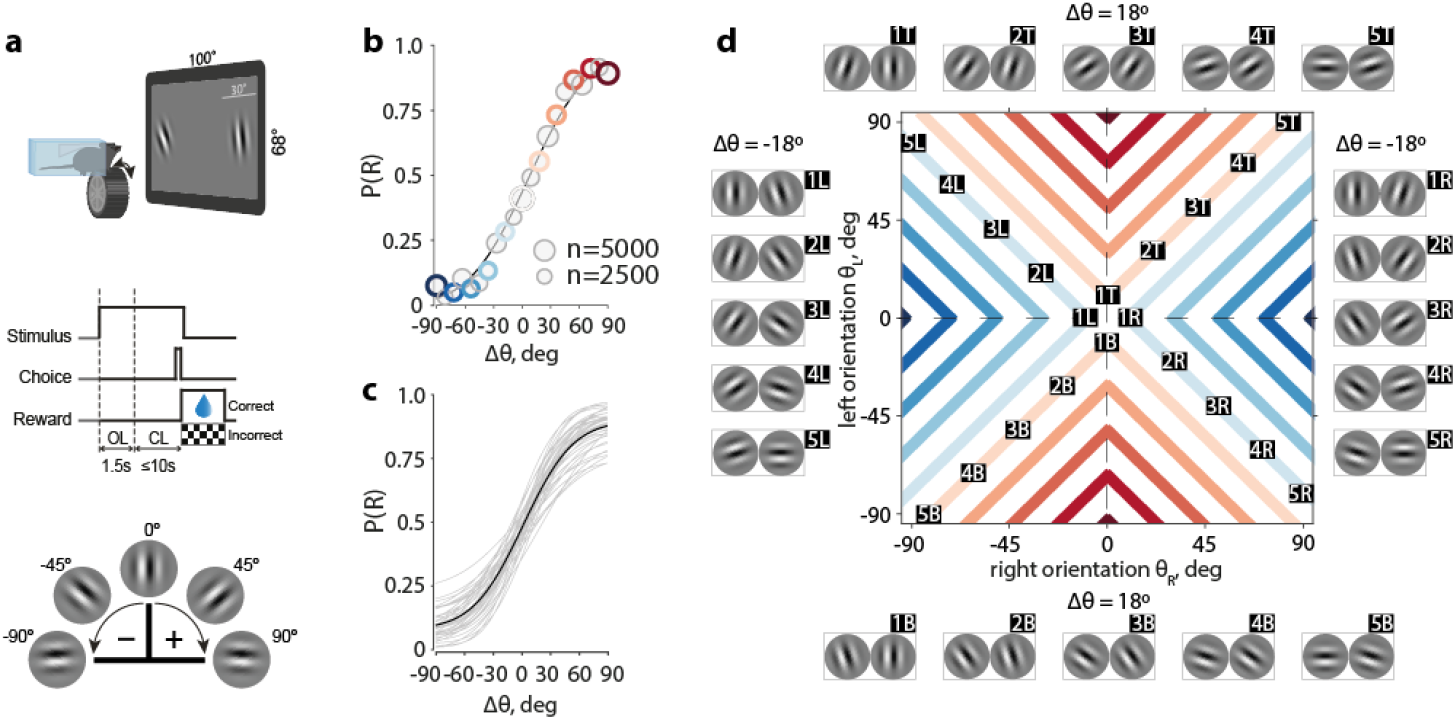
Mice successfully learned a novel invariant orientation discrimination task. **a.** Top. Schematic of a mouse during an experimental session. Middle. Epochs of one trial. OL – open loop, during which the wheel manipulator did not move the stimuli on the screen, CL – closed loop, during which it could. Bottom. Convention for the angle signs. **b.** Psychometric curve of an example animal. Solid line – best fit of the cumulative Gaussian psychometric function, circles – data points, circle sizes represent numbers of trials, colors correspond to colors in d, gray circles are data points not explicitly marked in d. **c.** Psychometric curves for all animals in the study, solid black line – population average. **d.** Many orientation pairs give the same task-relevant information quantified by angular separation or difficulty (Δ*θ*). Conditions with a fixed Δ*θ* in the 2d stimulus space (colored lines) correspond to Δ*θ* conditions (circles) of the same color in b. Example stimuli for four branches of constant Δ*θ* (two branches for 18°, and two for −18°) are displayed along the sides of the stimulus space map. Labels next to the images of orientation pairs correspond to labels on the stimulus space map.

#### Mice reach a high success rate in a relative orientation discrimination task

As an initial step in the analysis of choice behavior, we quantified performance as a function of task difficulty using a standard cumulative Gaussian psychometric function (Wichmann and Hill, 2001). We modelled the probability of choosing the right stimulus, P(R), as a function of the angular separation Δ*θ* = |*θ_L_*| – |*θ_R_*| between the two orientations, where |·| denotes the verticality, with small angular separations corresponding to difficult conditions and large angular separations corresponding to easy conditions. An angular separation Δ*θ* = 0 corresponds to two *equally vertical* orientations, which are not necessarily parallel. Conditions with Δ*θ* < 0 and Δ*θ* > 0 correspond to a more vertical orientation on the left and right side, respectively (example animal, **Fig. 1b**; population, **Fig. 1c**). The mice reached an average performance of 74.7 ± 0.7% correct, with an average sensitivity parameter of the psychometric curve *σ* = 42.93 ± 1.18°. Animals retained their performance level after introducing changes in spatial frequencies and stimulus sizes, suggesting their decision-making strategy did not rely upon these low-level statistical properties of the stimuli (average psychometric curves over the 3 sessions before and after changing either of these parameters did not differ from each other, **Supplementary Fig. 2**).

As this task disentangles any given probability of choice from specific orientations, a fixed difficulty Δ*θ* that corresponds to one point on the psychometric curve is given by many possible pairs of orientations (*θ_L_*, *θ_R_*). For example, Δ*θ* = 30° corresponds to orientation pairs (30°, 0°), (−60°, 30°), and many others (**Fig. 1d**). Conversely, no given orientation was always rewarded, since for any orientation (except 0°), there was a possibility that the other orientation was more vertical. For equally vertical orientation pairs, a side chosen at random was rewarded. Consequently, this task design compels the animal to estimate the *verticality* of the left and right orientations, |*θ_L_*| and |*θ_R_*|, and compare their estimates, rather than detect a learned orientation.

Animals may not strictly adhere to this ideal strategy, so long as they obtain sufficient amount of water reward in each experimental session. This amount can be difficult to estimate precisely, since it varies significantly from animal to animal, and depends on age, gender, food intake, and genetic background. We estimated that, with the choice variability taken into account, an animal looking at only one of the two stimuli will perform at 63.1 ± 0.6% correct, exceeding the 50% chance level, but, on average, not being able to maintain its weight at the pre-training level, assumed to be a heathy reference baseline. In the following section, we introduce a model that quantifies how animals combine information from the two orientations while also capturing deviations from the ideal strategy.

### Probabilistic choice model

#### Accounting for stimulus space and biases in the model

The psychometric curve quantifies an animal’s behavior along a single dimension of difficulty, Δ*θ.* However, given the task structure, the complete representation of the stimulus space is two-dimensional, with a unique stimulus condition corresponding to a pair of angles (*θ_L_,θ_R_*). In this space, a fixed Δ*θ* is given by all stimulus conditions along the iso-difficulty lines (*branches*) that lie in the four quadrants of the space and correspond to four different combinations of angle signs (**Fig. 1d**). We therefore considered the probability of choosing the right orientation, P(R), for all stimulus conditions in this space.

To get a better insight into the factors that affect the animals’ choices, we developed a psychometric model that provided a functional mapping from the two-dimensional stimulus space to the probability of the right choice, P(R). We assume that in every trial, a mouse makes noisy estimates 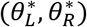 of both orientations (*θ_L_,θ_R_*), compares their verticalities 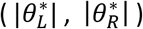, and makes a choice (**Fig. 2a**). The probability of a right choice P(R) in this procedure is expressed as an integral of the distribution of estimates 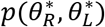 over the 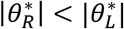 subspace (**Fig. 2b**) (Methods: Eqs. 1-4). The shape of the P(R) surface over the stimulus space (*θ_L_, θ_R_*) (**Fig. 2c**, left) is therefore determined only by the parameters of the distribution 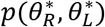, which we represent as a product of animal’s likelihood function over percepts and its prior distribution.

**Figure 2.**
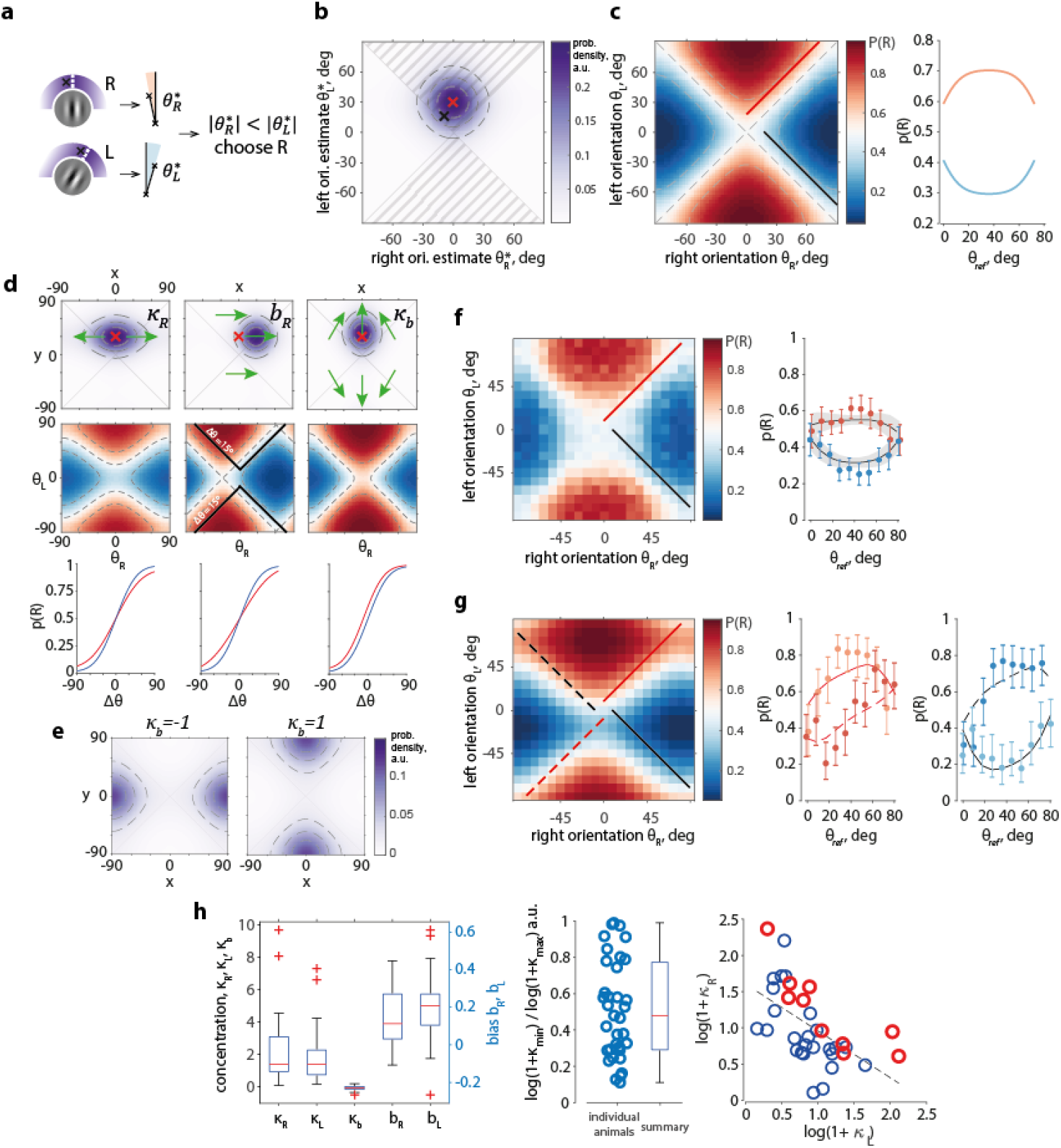
The choice model characterizes individual biases and strategies, and predicts variation of performance with reference orientation, as found in the data. **a.** Choice model schematic. The angles of two oriented Gabor patches (white dashed lines, left column) are estimated as samples from circular distributions (density – in purple, estimates – black crosses), their absolute values are measured as angular distances to the vertical and compared between each other (middle column), which generates a choice. **b**. Distribution 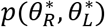 of orientation estimates as in a, in 2d space, for (*θ_L_, θ_R_*) = (30°, 0°) (red cross) in an unbiased model with (*κ_R_, κ_L_*) = (*2, 2*), and a sample from this distribution (black cross). Probability mass inside the shaded areas 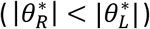 is equal to the probability of right choice P(R). Dashed lines – distribution quartiles. **c**. Left. P(R) of model in b evaluated at all stimulus pairs (*θ_L_, θ_R_*). Red and black lines – one example branch of Δ*θ* = 18° and Δ*θ* = –18° respectively. Right. P(R) along the branches of constant Δ*θ* marked on the left panel with a red and black lines. **d**. Effect of model parameters on animals’ likelihood distributions over percepts (top row), P(R) surface assuming uniform priors (middle row), and the corresponding psychometric curves (bottom row); red crosses – distribution means before parameter manipulation, green arrows – transformation of the distributions with parameter change; blue psychometric curves – before parameter change, red curves – after. The center panel in middle column shows P(R) values displaced relative to Δ*θ* isolines (solid black: example isoline for Δ0=15°). **e.** Choice priors *p_b_*(*x,y*) with *κ_b_* equal to −1 (left panel; left choice bias) and 1 (right panel; right choice bias). **f**. Left. Population average P(R) (n = 40 mice) with one example branch of Δ*θ* = 9° and Δ*θ* = –9° marked with red and black lines. Right: P(R) values as on the left panel (dots with error bars, mean ± c.i.), and average of model predictions (black lines with shaded areas, mean ± c.i.) across all animals. See Supplementary Fig. 3a for P(R) of every animal. **g**. Example mouse, left: P(R) of the fitted model, middle: P(R) along the red dashed and solid lines on the left panel predicted by the model (lines) and computed from the data (dots with error bars, darker dots correspond to the dashed line), right: P(R) along the black dashed and solid lines on the left panel, as predicted by the model (lines) and computed from the data (dots with error bars, darker dots correspond to the dashed line). See Supplementary Fig. 4d for the model of P(R) for every animal. **h**. Left: population summary of model parameters fitted to all mice (n=35; n=5 animals with *κ_R_* or *κ_L_* estimated on the edge of the allowed range of values are excluded). Middle: ratio of log(*κ_R_* + 1) and log(*κ_R_* + 1) with the smaller of the two values divided by the larger value for each mouse (n=35). Circles – individual animals. Box plot – population summary: red line – median value, box borders – 25^th^ and 75^th^ percentiles, whiskers are up to most extreme parameter values, red crosses – outliers. Right: log(*κ_R_* + 1) and log(*κ_R_* + 1) across the population are significantly anti-correlated. Linear regression line for all animals together. Red circles – 10 mice with best performance.

We model the likelihood *p*(*x,y*) as a product of circular von Mises functions 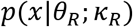 and *p*(*y*|*θ_L_*; *κ_L_*) centered at the values of *θ_R_* and *θ_L_* equal to the true orientations and with variability for each target quantified by the concentration parameters *κ_R_* and *κ_L_*. High concentrations correspond to low variability in the percepts, and *κ* is thus qualitatively inverse to the standard deviation and can be interpreted as the *certainty* (Drugowitsch et al., 2016; Laquitaine and Gardner, 2018). For example, a distribution of percepts *p*(*x,y*) is broader and shallower along the axis of lower concentration (**Fig. 2d,** left column, top), making P(R) more independent of the respective stimulus (**Fig. 2d,** left column, middle).

Percepts of each orientation can be systematically biased, with an animal consistently making choices as if the right or the left orientation were rotated more clockwise or counterclockwise. These systematic errors are accounted for by *translational biases b_R_* and *b_L_* (**Fig. 2d,** center column example: *b_R_* > 0, *b_L_* = 0), which move *p*(*x,y*) and consequently the P(R) surface relative to the angle axes without changing their values.

Both the translational biases and the certainty parameters change the slope of the psychometric curve but not its left-right choice bias (**Fig. 2d,** bottom row), with the effects generally indistinguishable in the Δ*θ* space as opposed to the complete stimulus space. A lower or higher certainty results in a shallower or steeper P(R) respectively, and a shallower or a steeper psychometric curve. On the other hand, a translational bias displaces the entire P(R) surface, overall decreasing performance for every Δ*θ* in the space of the psychometric curve.

To model a *choice bias* towards the right or left, we introduced a family of prior distribution functions or *choice priors p_b_*(*x,y*; *κ_b_*) parameterized by prior concentrations *κ_b_* (**Fig. 2e**). These choice priors cause an orientation on the right or on the left to effectively appear more vertical—as opposed to more clockwise or counterclockwise—or equivalently make an animal more certain about the verticality of that stimulus, or can be associated with procedural factors that similarly bias choices (**Fig. 2d**, right column) (Methods, Eq.2). For example, the choice prior for a rightward bias has a peak at (90°, 0°) (**Fig. 2e**, right, *κ_b_* > 0) and increases the probability of a right choice for any pair of orientations (**Fig. 2d**, right column) by biasing 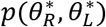 to the 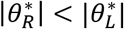 region (**Fig. 2d**, right column, green arrows).

Concentrations, translational biases, and a prior concentration {*κ_R_,κ_L_,b_R_,b_L_,κ_b_*} thus determine our model of choice, which allows a more complete analysis of P(R) than the psychometric curve. The model predicts a previously unexplored property of P(R): its variation along the branches of a fixed Δ*θ*. A model with zero biases and an equal certainty for both orientations (*κ_R_* = *κ_L_*) predicts a decrease in P(R) whenever either orientation is close to 0° or 90°, and an increase when close to 45 ° (**Fig. 2b-c**). We parameterized this variation using the *reference orientation θ_ref_* = min(|*θ_L_*|, |*θ_R_*|), i.e., the orientation of the more vertical stimulus. The source of this variation is clear from the position of 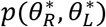 relative to the category boundary 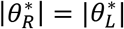 when considered along one branch of a fixed Δ*θ* (**Fig. 2b**): the probability mass of orientation estimates that result in error judgments (e.g., 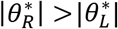 when |*θ_R_*| < |*θ_L_*|) is higher around *θ_ref_* = 0° and *θ_ref_* = 90° than around *θ_ref_* = 45°. This effect arises from the variability in both orientation estimates and their interaction with the category boundary in the circular space and cannot be replicated by the psychometric curve whose only input variable is Δ*θ.*

In summary, by combining information from two orientations, our model predicts a dependency of probability of choice not only on difficulty but also on reference orientation. This latter variability necessarily follows from the circularity in the input stimulus space given a limited certainty in orientation estimates.

#### The model captures the animals’ choices

We next analyzed the choices of the mice in the two-dimensional stimulus space. For the population of animals, P(R) varied with difficulty Δ*θ,* as expected from the psychometric curves (**Fig. 1b-c**), and with the reference *θ_ref_* (**Fig. 2f**), as predicted by our model (**Fig. 2b-c**). For a fixed Δ*θ* > 0, P(R) was higher (and choices were more often correct) when the orientations were far from horizontal or vertical (**Fig. 2f**), while for Δ*θ* < 0 P(R) was smaller (and the choices were also more often correct) when the orientations were far from horizontal or vertical.

The model reproduced this performance variation for individual animals (**Fig. 2g**). However, due to individual biases, the P(R) curves for fixed Δ*θ* were more distorted than in the unbiased case (cf. **Fig. 2c**, right). Counterintuitively, P(R) for the same Δ*θ* in different quadrants of the stimulus space could represent on average opposite choices (**Fig. 2g**, center, right), which our model accounted for thanks to translational biases. The model successfully captured animal-specific differences in choice probabilities (**Supplementary Fig. 3**), explained the data significantly better than the psychometric curve (ΔAIC = 798.8 ± 141.9; ΔAIC > 0 for all animals), and explained significantly more deviance (Runyan et al., 2017) (ΔFDE = 9.03 ± 1.48%, p = 1.07 · 10^-6^, signed-rank test).

Across the population of animals, the average stimulus concentration values were high and positive (*κ_R_* = 2.22 ± 0.69, p = 3.73 · 10^-7^; *κ_L_* = 1.76 ± 0.52, p = 1.34 · 10^-7^, t-tests) (**Fig. 2h**, left) showing that the animals used both targets for the decision. The bias concentration *κ_b_* was small (*κ_b_* = −0.06 ± 0.05, p = 0.01), indicating a mixed bias across the population. The translational biases (*b_R_* = 0.14 ± 0.05, p = 3.71 · 10^-6^; *b_L_* = 0.19 ± 0.05, p = 1.21 · 10^-7^) were similarly small but significant.

Although the stimulus protocol, reward sizes, and session schedules were designed to motivate animals to use information about both orientations equally, we found that the strategies of individual animals ranged from a balanced orientation comparison to a reliance on one target more than the other. We quantified this range of strategies with the ratio of the log of the concentrations *κ_R_* and *κ_L_*, with ratios closer to 1 representing more balanced strategies (**Fig. 2h**, center). The right and left concentrations were significantly anti-correlated (ρ = −0.57, p = 4.45 · 10^-4^; t-test, criterion α = 5 · 10^-3^ corrected for multiple comparisons), reflecting a trade-off in animals that preferentially used information from one of the stimuli (**Fig. 2h**, right). Despite this trade-off, the best-performing animals also had higher concentrations overall (p < 0.05; ANCOVA, F-test of intercept with fixed slope), showing that while the task permitted relative flexibility in choice strategies, a more accurate estimation of the orientations was necessary to achieve a high success rate. Other parameters of the model did not significantly correlate with each other or with the concentration ratios.

In summary, our model accounted for biases in the animals’ behavior and explained the performance variation with *θ_ref_*. Individual animals weighted sensory information from the two orientations differently, following strategies that were *sufficient* to obtain needed amounts of reward, but were not perfectly aligned with the true stimulus-reward space. While left and right concentrations were anticorrelated across the population, high success rates required overall high certainty in the orientation estimates.

#### Discrimination acuity

We next used our model to estimate the minimal orientation difference the animals could reliably discriminate. A change in a pair of orientations that results in a significant change in P(R) is the smallest for conditions with the largest gradient of P(R). Since the numerical gradient directly computed from the data can be too noisy, we used our model to more accurately find the maximum gradient conditions. After identifying these conditions, we used experimentally obtained trial outcomes to test the significance of P(R) change.

Following this procedure, we compared probability of a right choice in stimulus conditions with the highest gradient and in neighboring conditions (**Fig. 3**). We found that a change in either left or right orientation by 9° resulted in a significant change in P(R) for 62.5% (n = 25) of animals, and that a change by 27° resulted in a significant change for all (n = 40) animals (**Fig. 3a-e**). For the only animal tested with a 3° sampling of stimuli, we found that changing both orientations by 3° along or against the gradient—amounting to a total change of 6°—resulted in a significantly different P(R) (p < 0.0005, both cases) (**Fig. 3f-i**).

**Figure 3.**
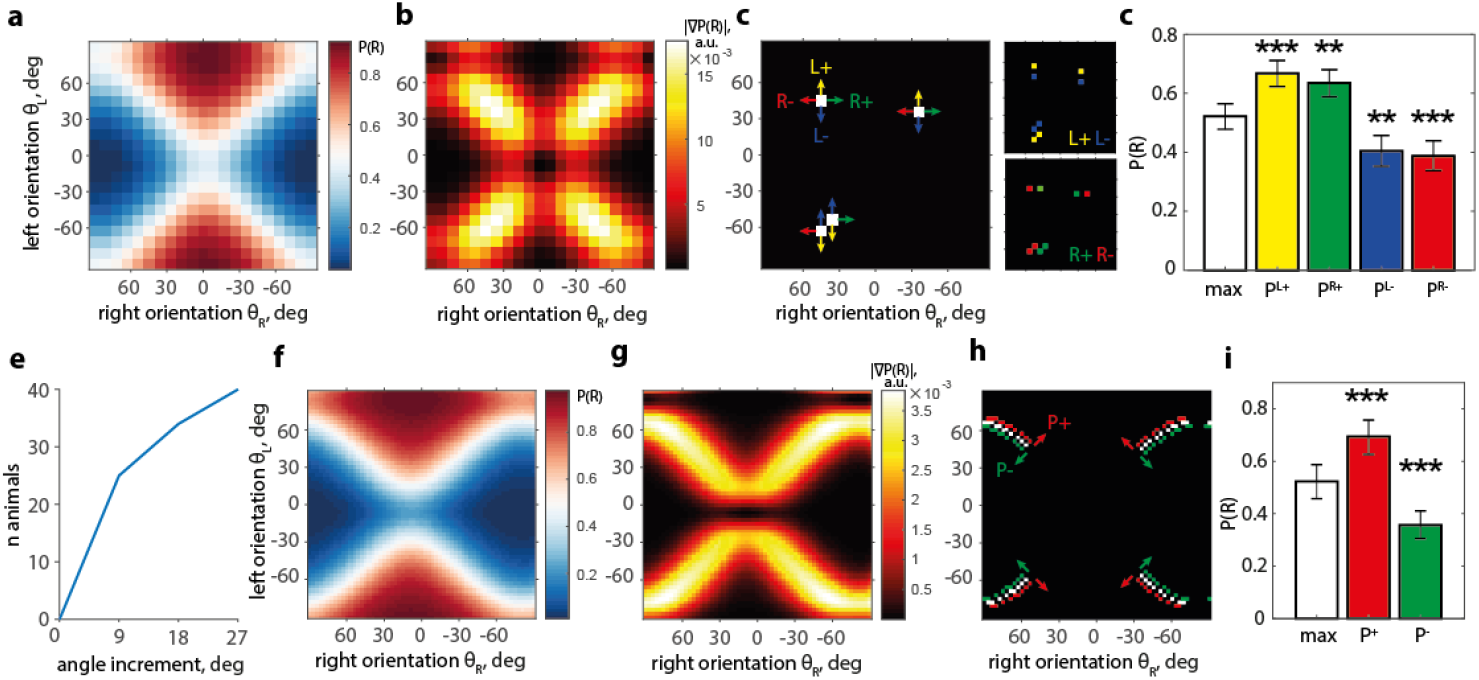
Mice can reach an orientation acuity of 6°. **a**. P(R) model surface of an example mouse. **b**. |VP(R)| absolute value of the gradient of P(R) surface in a. **c**. Four stimulus conditions (white) with | VP(R) | in the top 5% of values and P(R) close to 0.5 (0.48 < P(R) < 0.52). Arrows of the same color show angle change yielding neighboring conditions: P^L+^ (yellow; “L+” for left stim. change that increases P(R)), P^R+^ (green), P^L-^ (blue), P^R-^ (red). Insets (right) show these conditions in the stimulus space. **d**. Pooled P(R) in maximum gradient conditions (white), P^0.5^ = 0.52 ± 0.04, differs from P(R) in the four neighboring conditions P^L+^ = 0.67 ± 0.04, p < 2.5 · 10^-4^ (yellow), P^R+^ = 0.63 ± 0.05, p < 2.5 · 10^-3^ (green), P^L-^ = 0.40 ± 0.05, p < 2.5 · 10^-3^ (blue), P^R-^ = 0.39 ± 0.05, p < 2.5 · 10^-4^ (red) (binomial confidence intervals, χ^2^ test, df = 1, n = 4 comparisons). **e.** Cumulative number of animals for which at least one direction of angle change gives a P(R) significantly different from P^0.5^, as a function of angle change. **f,g**. Similar to a,b for an animal trained with 3° angle binning. **h.** Maximum gradient conditions (white, same criteria as in c), and neighboring conditions obtained by changing both angles by ±3° in the direction of P(R) increase (P^+^, red), and decrease (P^-^, green). **i.** Pooled P(R) in three groups highlighted in h: P^0.5^ = 0.52 ± 0.04, P^+^ = 0.69 ± 0.07 (red), P^-^ = 0.36 ± 0.05 (green), both different from P^0.5^ with p < 0.0005 (χ^2^ test, df = 1; n = 2 comparisons).

In summary, our model allowed an in-depth analysis of discrimination acuity by utilizing a complete picture of the P(R) gradient and identifying stimulus conditions where the sensitivity to angle change was the highest. We found that an angle change of 6° can be significantly detected based on the change of probability of choice, thus establishing a lower bound for mouse orientation discrimination acuity.

#### Effects of trial history

Choice strategies are determined not only by preferential weighting of available sensory information but also by trial history (Abrahamyan et al., 2016; Akrami et al., 2018; Busse et al., 2011; Corrado et al., 2005; Fründ et al., 2014; Urai et al., 2017; Yu and Cohen, 2008). To account for history-related biases, we included a history prior *p_h_*(*x,y*) parameterized with a concentration parameter *κ_h_* and a term *h* that linearly depended on the choice *r* and target orientation *s* in the previous trial through history weights (Busse et al., 2011; Corrado et al., 2005; Fründ et al., 2014), *h* = *sh_s_* + *rh_r_*. A pair of weights (*h_s_,h_r_*) determined the choice strategy of an animal, such as “win-stay” (**Fig. 4a**, model example) or “lose-stay” (**Fig. 4b**, example animal) throughout all trials, and in combination with the choice and target of the previous trial (r, s) resulted in the history-dependent change of the P(R) (**Supplementary Fig. 4a-e**) and the psychometric curve (Fründ et al., 2014) (**Fig. 4a,b**).

**Figure 4.**
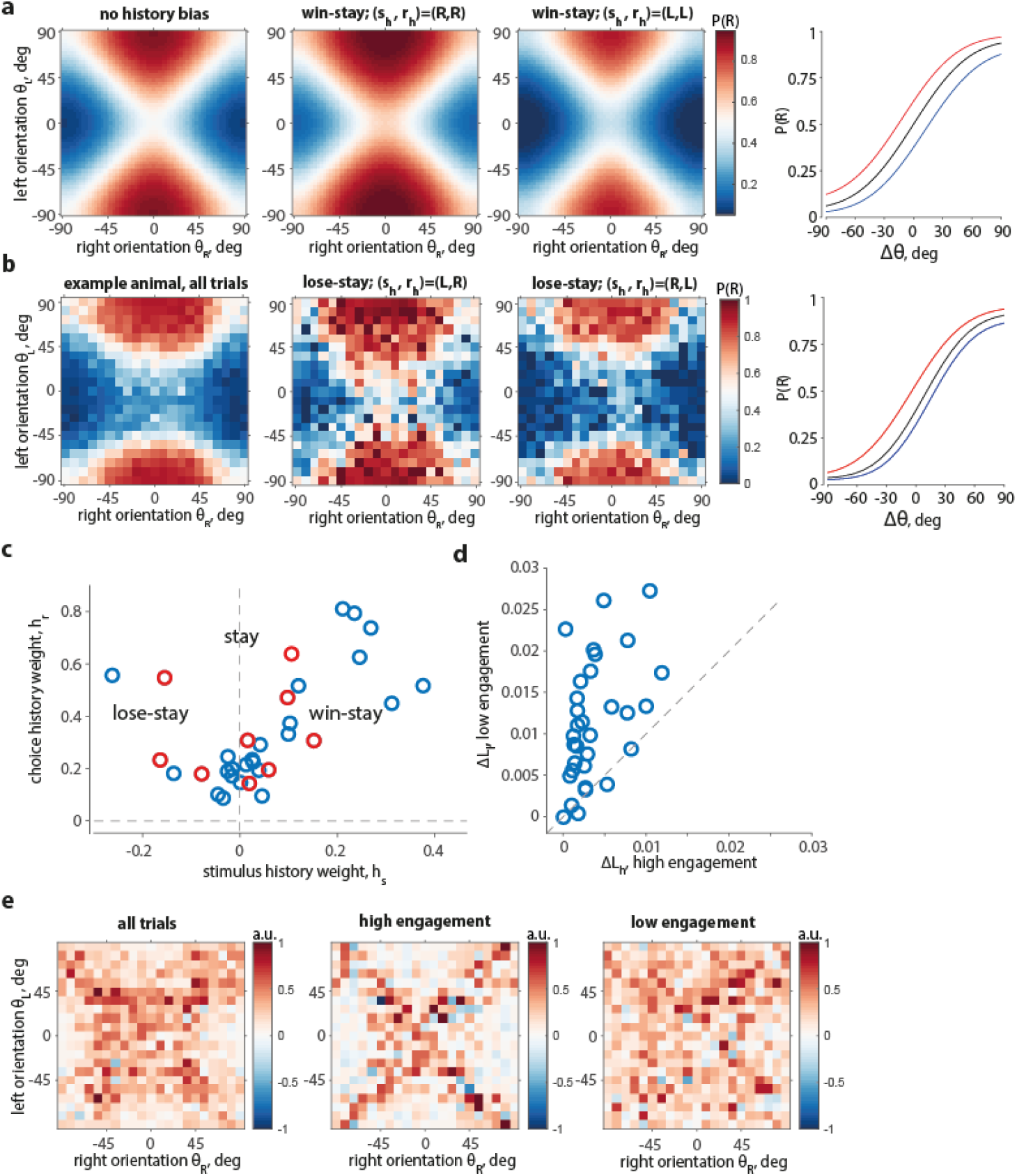
The probability of choice is affected by the choice and reward on the previous trial, with a larger effect during periods of lower task engagement. **a**. P(R) of an unbiased example model (*κ_R_,κ_L_,κ_b_,b_R_,b_L_*) = (1.5, 1.5, 0, 0, 0) with a win-stay strategy (*h_s_, h_r_, κ_p_*) = (0.4, 0.4, 0.5); from left to right: [1] without history bias (after a ‘neutral’ trial, (*s_h_,r_h_*) = (0,0)), [2] after a successful right choice (*s_h_,r_h_*) = (1,1) with P(R) biased to the right choices as a result, [3] after a successful left choice (*s_h_,r_h_*) = (–1, –1) with P(R) biased to the left choices, [4] psychometric curves corresponding to [1-3]: without a history effect (black), after a correct right choice (red), after a correct left choice (blue). **b**. P(R) of an example animal with large history biases (*h_s_, h_r_*) = (–0.26, 0.55) (lose-stay strategy); from left to right: [1] average P(R) on all trials, [2] P(R) after an unsuccessful right choice (*s_h_,r_h_*) = (–1,1) biased to right choices, [3] P(R) after an unsuccessful left choice (*s_h_,r_h_*) = (1,–1) biased to left choices; P(R) of all conditions in [2] and [3] with fewer than N = 5 trials was set to the average P(R) of its neighbors; [2] and [3] are conditioned on preceding errors and thus had a relatively low number of trials since the performance of all mice exceeded a 65% success rate, [4] psychometric curves of the corresponding surfaces, colors as in a. **c**. History weights (*h_s_, h_r_*) and corresponding strategies of all animals (4 outliers not shown); “stay” and “win-stay” strategies are predominant, with few examples of “lose-stay”. Blue circles – animals trained to detect a more vertical orientation, red circles – more horizontal orientation. **d**. Increase in the log-likelihood due to inclusion of history terms is larger for low-engagement trials than high-engagement trials. Abscissa – difference (Δ*L_h_*) between average log-likelihood under the model with history (*p_h_*) and without history (*p*_0_) of high-engagement trial outcomes (*r^h^*), Δ*L_h_* = 〈*L*(*r^h^,p_h_*)〉 – 〈*L*(*r^h^,p*_0_)〉, ordinate – difference (Δ*L_l_*) between average log-likelihood under the model with and without history of the low-engagement trial outcomes (*y_l_*), Δ*L_l_* = 〈*L*(*r^1^, p_h_*)〉 – 〈*L*(*r^1^, p*_0_)〉. **e**. Increase in the log-likelihood due to inclusion of history terms changes with stimulus conditions and engagement modes; left to right: [1] Δ*L_θ_* – average log-likelihood difference for every stimulus condition, Δ*L_θ_* = 〈*L*(*r,p_h_*)〉_*θ*_ – 〈*L*(*r,p*_0_)〉_*θ*_, all trials are taken, maps are Z-scored and averaged across animals, conditions with fewer than 10 trials are excluded, [2] Δ*L_hθ_* – same value computed for high-engagement trials only, [3] Δ*Lι_θ_* – same value computed for low-engagement trials only.

Through the flexible family of history priors, our model captured a variety of strategies in addition to win-stay (**Supplementary Fig. 4f**). Most of our mice showed a mild tendency for the “stay” strategy, followed by the “win-stay”, and, rarely, the “lose-stay” strategy (**Fig. 4c**), largely in consistency with the previous report (Odoemene et al., 2018). The history-dependent model explained the data significantly better than the history-independent model (ΔAIC = 211.8 ± 40.1; ΔAIC > 0 for all but n = 4 animals), and explained significantly more deviance (ΔFDE = 5.07 ± 0.98 %, p = 8.1 · 10^-6^, signed-rank test).

We investigated whether the animals relied on history to a different extent during periods of relatively high and low engagement in the task, which we identified based on performance within a session. Performance fluctuations correlated with changes in biomarkers typically associated with task engagement and overall attentiveness to the task (McGinley et al., 2015; Reimer et al., 2016) (shown using the same behavioral protocol in Abdolrahmani et al., 2021) (Abdolrahmani et al., 2021). For each of the two engagement levels, we computed the difference between the trial-average log-likelihood of choices given a model with a history prior and without: Δ*L_l_* for low-engagement trials, and Δ*L_h_* for high-engagement trials. The increase in explanatory power was larger for the low-engagement trials (Δ*L_l_* > Δ*L_h_*) (Wilcoxon test, p=2.12 · 10^-7^) (**Fig. 4d**), which indicated that during the low engagement trials, the choices were more strongly driven by the history biases.

Difficult stimulus conditions were more susceptible to the influence of history priors than easy conditions (**Fig. 4e**, left), in a way that depended on the engagement state of the animal. During periods of high engagement, the inclusion of history priors led to a substantial improvement in the model performance only for the most difficult conditions (**Fig. 4e**, center), while in the low-engagement periods, most stimulus conditions were affected (**Fig. 4e**, right).

In summary, after expanding our model to capture history-dependent biases, we found that the most prominent strategies were “win-stay” and “stay”, and that choices were affected by history biases to a greater extent during periods of lower engagement. Our observations demonstrate that choice heuristics can fluctuate together with the cognitive state of the subject.

## Discussion

Using high-throughput automated cages with voluntary head fixation, we trained a large cohort of mice (n = 40; 1,313,355 trials) in a complex variant of a 2AFC orientation discrimination task. The task required the mice to measure the relative orientations of two stimuli, thereby decoupling choice from the particular orientation of an individual stimulus. We quantified their behavior with a novel model of choice that accounted for the circularity of the stimulus space and for individual choice biases and strategies. The model explained variation in the probability of choice not only with the task difficulty Δ*θ*, but also with the reference orientation *θ_ref_*, an effect not reported previously. With the help of the model we found that the maximum acuity of orientation discrimination in expert animals can be as small as 6°. Our model could be easily extended to examine history biases, ubiquitous in human and animal psychophysical experiments (Abrahamyan et al., 2016; Akrami et al., 2018; Busse et al., 2011; Corrado et al., 2005; Fründ et al., 2014; Urai et al., 2017; Yu and Cohen, 2008), revealing a modulation of history components by the animals’ engagement, affecting choices more strongly and over a broader set of stimulus conditions whenever the engagement was relatively low. Our work responds to the need for a visual task that depends on abstract choice categories and is invariant to specific visual stimuli, but can be learned by mice, relies on basic visual features, and allows straightforward quantification within the probabilistic modelling framework. We argue that in addition to these advantages, our task can be useful in engaging higher visual areas in the computation of decision (DiCarlo and Cox, 2007), and can provide valuable insight into the relationship between neural and behavioral variability (Beck et al., 2012; Britten et al., 1996; Brunton et al., 2013; Drugowitsch et al., 2016; Renart and Machens, 2014).

### Behavioral assays for studies of perceptual invariances and their quantifications

Our task will be particularly advantageous for the study of the neural mechanisms underlying perceptual invariances. With the availability of unique experimental toolboxes, the mouse is currently the animal model of choice for the dissection of neural circuits (Abbott et al., 2020; Luo et al., 2018; Madisen et al., 2015). However, although visual behaviors elicited by low-level visual features have been well characterized (Huberman and Niell, 2011; Zoccolan et al., 2015), intermediate (e.g. textures) and high-level vision (e.g. objects) are largely unexplored in this species. Therefore, mouse studies that utilize complex visual stimuli are challenged by (1) the well-known difficulty of parameterizing complex objects (DiCarlo and Cox, 2007; DiCarlo et al., 2012; Riesenhuber and Poggio, 2000), (2) the unknown neural substrate that encodes these parameters, (3) the uncertainty about whether mice can learn the task in a reasonable time—if at all—and (4) the difficulty in inferring behavioral strategies given the parametric complexity of the stimulus space (Alemi-Neissi et al., 2013; Vermaercke and Op de Beeck, 2012). Our task represents a convenient solution: it builds upon existing orientation discrimination tasks in mice (Andermann et al., 2010; Goard et al., 2016; Long et al., 2015; Reuter, 1987; You and Mysore, 2020), in which a specific orientation is to be chosen over a distractor orientation (Andermann et al., 2010; Long et al., 2015; Pinto et al., 2013; Poort et al., 2015; Resulaj et al., 2018; Reuter, 1987; You and Mysore, 2020), or in which a change relative to a specific orientation is to be detected (Glickfeld et al., 2013; Jin et al., 2019; Wang et al., 2018, 2020). However, it complexifies the discrimination by introducing well-controlled invariances (to specific orientations, spatial frequency, and stimulus size), exploring stimulus dimensions that are easy to parameterize and that have a clear neural representation, and can be learned by mice in a reasonable time.

Our model helped estimate orientation discrimination acuity, which reached a 6° angle difference for one mouse tested with the smallest angular separation of 3°. Constrained by limitations related to a different study, we did not attempt to train animals at smaller angular differences, so it is possible mice can discriminate orientation differences even smaller than 6°. The orientation discrimination acuity of mice has been previously measured in a 2AFC tasks with a distractor (Reuter, 1987), and change detection tasks (Glickfeld et al., 2013; Jin et al., 2019; Wang et al., 2018, 2020). Acuity measures have been reported as thresholds or just-noticeable differences (JNDs) and commonly rely on model-derived values, such as the model-based inverse of a certain success rate (Glickfeld et al., 2013; Jin et al., 2019), the mean of the fitted Gaussian (Wang et al., 2018, 2020), or 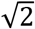 times its standard deviation (Wang et al., 2018, 2020). We developed a new acuity estimation procedure suitable for our stimulus space, in which we identified stimulus conditions with the highest gradient of model-predicted P(R) and compared the performance in these and neighboring conditions. Our approach took advantage of the complete stimulus space representation of P(R) instead of relying on a cruder psychometric model to compute a JND or a threshold value.

### Task- and behavior-related factors influencing choice

By parameterizing biases, history effects, and orientation certainties, our model showed that animals largely followed the intended choice strategy, but also exhibited variation that could be interpreted as animal-specific choice heuristics. One such heuristic was evident in the trade-off of concentration values, with some animals unequally weighting stimulus information. Accuracy of orientation estimation was still necessary for high success rates, but even among the best-performing animals right and left concentrations were anti-correlated. This trade-off demonstrated that animals followed a range of “sufficiently good” strategies when solving the discrimination problem.

Such strategies can be interpreted as examples of suboptimal or approximate inference in an uncertain environment. Suboptimal inference is sometimes thought of as an adaptive phenomenon, a way for a subject to deal with the complexity of the task at hand by constructing and acting upon its approximate model (Beck et al., 2012). Adherence to a suboptimal strategy can be linked to limited cognitive resources (Whiteley and Sahani, 2012; Wyart and Koechlin, 2016), which in our task fluctuate together with task engagement. Indeed, we find that history-dependent biases—another manifestation of suboptimal behavior—are stronger during periods of lower engagement. We demonstrate this by introducing history priors—in a form that allows their analytical inclusion into our model—that increase the explanatory power of the model more in periods of lower engagement than in periods of higher engagement. These fluctuations of the history biases are driven by the internal state of the animal, are independent of the stimulus protocol, and thus will occur in addition to difficulty- or confidence-dependent fluctuations, as recently described (Lak et al., 2020). During periods of decreased performance, higher explanatory power of history terms is not guaranteed, but it is consistent with switching between history-driven and stimulus-driven choice modes (Ashwood et al., 2020).

### Limitations of our approach

Although we believe that our work substantially advances the understanding of mouse behavior during complex orientation discrimination, our approach has limitations at the level of model design and strategy interpretation. First, our model assumes fixed psychometric parameters across sessions and trials, and thus a more flexible, dynamically parameterized model could give a better insight into biases and choice strategies of mice. Second, the goodness of fit of the model with respect to the variation of P(R) with *θ_ref_* could be further improved: in some animals this variation is larger than the model prediction (**Supplementary Fig. 5**, example animal), which could be explained by a dependency of *κ_R_* and *κ_L_* on the proximity to the category boundary 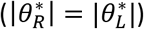 (Jazayeri and Movshon, 2007). Finally, direct interpretation of concentration values might not be directly relatable to perceptual sensitivity, since they were likely decreased by nonsensory factors, such as noise in the decision computation (Beck et al., 2012; Dosher and Lu, 1998; Drugowitsch et al., 2016), inherent priors (Girshick et al., 2011), and choice heuristics (Beck et al., 2012; Gardner, 2019).

### Future directions and potential implications

Since our task relies on perceptual invariances and decouples the decision information from specific sensory stimuli, it can be useful for exploring the neural basis of decision-making in the future studies. A similar task design relying on combinations of stimuli has been used extensively in the decision-making literature (Constantinople et al., 2019; Hernández et al., 1997; Jogan and Stocker, 2014; Pinto et al., 2018; Scott et al., 2015; Steinmetz et al., 2019), but has not been reported in mouse orientation discrimination experiments.

Furthermore, our task can give a valuable insight into the relationship between neural and behavioral variability. Whether behavioral variability arises predominantly from sensory sources (Brunton et al., 2013), or from the deterministic or stochastic suboptimality of decision computation (Beck et al., 2012) is one of the central questions in the neuroscience of decision-making. The complexity of our orientation discrimination task will increase the role of suboptimal decision computation, as has been predicted theoretically (Beck et al., 2012; Gardner, 2019; Whiteley and Sahani, 2012), and will provide an opportunity to study the correlates of this suboptimality in the neural responses.

Finally, our task is well suited for isolating the contributions of visual cortical areas in the computation of decision. The importance of a particular visual area for decision-making depends on the type of task (Pinto et al., 2019): mice with a lesioned or silenced visual cortex have shown above-chance performance in detection paradigms (Glickfeld et al., 2013; Prusky and Douglas, 2004), possibly reflecting a predominant role of the superior colliculus (Wang et al., 2020), while for orientation discrimination tasks with a distractor, the visual cortex is necessary (Jurjut et al., 2017; Poort et al., 2015; Resulaj et al., 2018). Our incrementally more complex version of the orientation discrimination task could provide further insight into the role of V1 and downstream visual areas in the computation of decision (DiCarlo and Cox, 2007), and thus could be a useful addition to the common behavioral protocols for mice.

## Acknowledgments

We thank Yuki Goya and Rie Nishiyama for their support with behavioral training. We thank O’Hara and CO., LTD., for their support with the equipment.

This work was funded by RIKEN BSI and RIKEN CBS institutional funding, JSPS grants 26290011, 17H06037, C0219129 to AB, and Fujitsu collaborative grant.

## Methods

### Experimental Model and Subject Details

All surgical and experimental procedures were approved by the Support Unit for Animal Resources Development of RIKEN CBS. We used n = 40 transgenic mice: Thy1-GCaMP6f (n = 37), Camk2-tTA TRE-GCaMP6s (n = 2), Emx1-tTA TRE-GCaMP6s (n = 1), with a total of 30 male and 10 female animals, aged 4 to 25 months. The triple transgenic strain Camk2-tTA TRE-GCaMP6s was established by cross-mating Camk2a-cre and Camk2a-tTA. The triple transgenic strain Emx1-tTA TRE-GCaMP6s was established by cross-mating Emx1-cre and Camk2a-tTA.

Animals were anesthetized with gas anesthesia (Isoflurane 1.5-2.5%; Pfizer) and injected with an antibiotic (Baytril®, 0.5 ml, 2%; Bayer Yakuhin), a steroidal anti-inflammatory drug (Dexamethasone; Kyoritsu Seiyaku), an anti-edema agent (Glyceol®, 100 μl, Chugai Pharmaceutical) to reduce swelling of the brain, and a painkiller (Lepetan®, Otsuka Pharmaceutical). The scalp and periosteum were retracted, exposing the skull, then a 4 mm-diameter trephination was made with a micro drill (Meisinger LLC). A 4 mm coverslip (120~170 μm thickness) was positioned in the center of the craniotomy in direct contact with the brain, topped by a 6 mm diameter coverslip with the same thickness. When needed, Gelfoam® (Pfizer) was applied around the 4 mm coverslip to stop any bleeding. The 6 mm coverslip was fixed to the bone with cyanoacrylic glue (Aron Alpha®, Toagosei). A round metal chamber (6.1 mm diameter) combined with a head-post was centered on the craniotomy and cemented to the bone with dental adhesive (Super-Bond C&B®, Sun Medical), mixed to a black dye for improved light absorbance during imaging.

After the implantation of the head-post and recovery from the surgery, for 2 weeks mice were placed in habituation cages with enriched environment, where they learned to obtain water from an apparatus similar to the automatically latching part of the behavioural setup. Next, mice were placed under a water restriction plan for 2 weeks, obtaining 3 ml of water a day during the first week, and 2 ml during the second, with a target of maintaining their body weight at 75-80% of the initial weight. If at this or any later point their weight dropped below the target level, mice were given additional water proportionate to the weight to be restored. After 2 weeks animals were moved to the training cages.

### Behavioral training

During training, animals were housed in individual cages connected to automated setups (Aoki et al., 2017) (O’Hara & CO., LTD., http://ohara-time.co.jp/) where two experimental sessions per animal per day were carried out. Sessions were initiated by animals themselves as they entered the setup and their head plate was automatically latched. Animals were trained in a 2AFC orientation discrimination task. Two oriented Gabor patches (20° visual angle static sinusoidal gratings, sf = 0.08 cpd, with randomized spatial phase, and windowed by a 2D Gaussian envelope with *4σ* equal to stimulus diameter) were shown on the left and right side of a screen positioned in front of the animal (LCD monitor, 25 cm distance from the animal, 33.6cm × 59.8 cm [~58° × 100°dva], 1080 x 1920 pixels, PROLITE B2776HDS-B1, IIYAMA) at ±35° eccentricity relative to the body’s midline. Mice reported which of the two stimuli was more vertical (more horizontal for n = 12 animals; task details in “Phases of training”) by rotating a rubber wheel with their front paws, which shifted the stimuli horizontally on the screen. For a response to be correct, the target stimulus had to be shifted to the center of the screen, upon which the animal was rewarded with 4 μL of water (amount adjusted for a few animals with non-typical weight and age). Incorrect responses were discouraged with a prolonged (10 s) inter-trial interval and a flickering checkerboard stimulus (2 Hz). If no response was made within 10 s (time-out trials), neither reward nor discouragement was given.

All trials consisted of an open-loop period (OL, 1.5 s) during which the wheel manipulator did not move the stimuli on the screen, and a closed-loop period (CL: 0—10 s) during which the wheel controlled their position. Inter-trial interval was randomized (ITI: 3— 5 s). Stimuli appeared on the screen at the beginning of the OL.

### Phases of training

Training in the automated behavioral setup went through three phases. First, the animal learned to rotate the wheel manipulator and was rewarded for consistent rotations to either side. During this phase no visual stimulus was presented. In the next phase, the animal was shown one vertical target (horizontal, n = 12), on one side of the screen chosen at random, and was rewarded for moving it into the center of the screen. In the final phase, the animal was shown two orientations, and had to move the more vertical (horizontal) one into the center of the screen. Since both stimuli moved synchronously with wheel rotation, the non-target stimulus moved out of the screen. In this phase, we sampled both orientations at random from a range of angles between −90° and 90°, with *θ >* 0 corresponding to clockwise and *θ* < 0 – to counter-clockwise orientations relative to the vertical (**Fig. 1a**). Orientations were initially sampled with a minimal angular difference of 30°, i.e. with specific angles from the set {-90°, −60°, −30°,0°,30°,60°} (−90° and 90° are the same orientation). As the animal’s performance reached 70% success rate on 5-10 consecutive days, we increased the difficulty by sampling angles at 15° angle difference, and later in the training – at 9°, with one animal’s conditions eventually sampled at 3°.

### Psychometric curve

We fitted the animal’s probability of making a right choice P(R) as a function of task difficulty using a psychometric function *ψ*(Δ*θ*; *α,β,γ,λ*) = *γ* + (1 –*γ* – *Λ*) *F*(Δ*θ*; *α,β*), where *F*(*x*) is a Gaussian cumulative probability function, *α* and *β* are the mean and standard deviation, *γ* and *λ* are left and right (L/R) lapse rates, Δ*θ* is the difference in the angular distance to the vertical, Δ*θ* = |*θ_L_*| – |*θ_R_*|. Confidence intervals were computed by bootstrapping (n = 999).

### Data selection

We analyzed trials from sessions in which the average success rate was at least 60%, and the proportion of time-out trials did not exceed 20%. We only used animals that had reached the minimal angular difference of 9°, and included the choice data from preceding sessions with minimal differences starting from 30°. We excluded the first trial of every session, all time-out trials and every trial that followed a time-out. The two dimensions of the stimulus space were flipped for horizontal-reporting animals when fitting our model. Same stimulus space transformation was done for all the population summaries where mice trained on horizontal targets were pooled together with mice trained on vertical targets.

### Model design

On each trial *i* the animal was shown a pair of stimuli {*θ_Ri_*, *θ_Li_*}, and made a right or a left choice *r_i_*, which we set by convention to be *r_i_* = 1 or = 0 respectively. We denote response and correct target on the previous trial as *r_hi_* and *s_hi_* respectively, with *r_hi_* = –1 or *r_hi_* = 1 if the animal chose left or right respectively, and *s_hi_* = −1 or *s_hi_* = 1 if the correct answer was respectively left or right, and *s_hi_* = 0 if targets had an equal verticality.

A choice in trial *i* was based on animal’s estimates 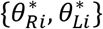 of the presented stimulus orientations 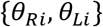. We model 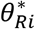 and 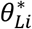 as random variables distributed according to a posterior distribution 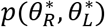 obtained after combining an animal’s likelihood distribution over percepts *p*(*x, y*) with prior terms *p_b_*(*x,y*) and *p_h_*(*x,y*) that model choice bias and history-dependent bias respectively. We reserve the (*x,y*) notation for the random variables modelling percepts and biases, and 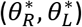 to refer specifically to the posterior over animal’s estimates, to which the decision rule is applied. We model the likelihood as a product of von Mises distributions *p*(*x*) and *p*(*y*) centered at *θ_Ri_* and *θ_Li_* respectively, with additional angle estimation biases (*translational biases*) *b_R_,b_L_* and with concentrations *κ_R_*, *κ_L_* (high concentration means smaller spread, with *κ* analogous to 1/*σ* of a normal distribution; only *κ* ≥ 0 were allowed) **(Fig. 2b,d)**:

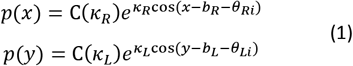

where *C*(*κ*) = 1/2*πI*_0_(*κ*), and *I*_0_ is modified Bessel function of order 0. A bias prior *p_b_*(*x,y*) that induces choice bias for right or left stimuli, and a history prior *p_h_*(*x, y*) that models choice dependency on previous choice and stimulus (*r_hi_* and *s_hi_*), are modeled as:

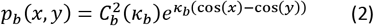

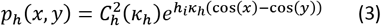

Here, *κ_b_* is a concentration parameter that regulates the strength and sign of choice bias, *κ_h_* is a concentration parameter of history prior, *h* = *h_s_s_hi_* + *h_r_r_hi_* determines the influence of the previous stimulus *s_hi_* and choice *r_hi_* with respective weights *h_s_* and *h_r_* fixed for a given animal, and *C_h_* = 1/2*πI*_0_(*κ_h_h_i_*) and *C_b_* = 1/2*πI*_0_(*κ_b_*) are normalization constants.

Since by convention we set vertical orientation to zero, the angle with the smaller absolute value is the correct choice. Hence, the probability of choosing right on a given trial is given by:

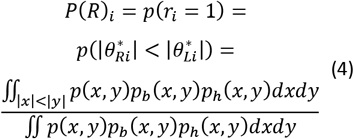

Overall, the model has eight fitted parameters (*h_r_,h_s_,κ_R_,κ_L_,κ_h_,b_R_,b_L_,κ_b_*), or five parameters (*κ_R_, κ_L_, b_R_, b_L_, κ_b_*) when we fit a history-free model. All angles were converted from (−90°, 90°) range to (180°, 180°) to satisfy periodicity.

Our model design follows similar models of perceptual inference(Girshick et al., 2011; Laquitaine and Gardner, 2018; Stocker and Simoncelli, 2006) with two distinctions. First, since our animals never report point estimates of the observed orientations—usually modeled as maximum a posteriori (MAP)—estimates only enter our model as not directly observed random variables. Second, since all orientations in our study are presented at 100% contrast, without added noise or any other form of stochasticity, and are displayed for the full duration of the trial (11.5 s or less if the choice is made earlier), we assume that the sensory evidence given by a specific orientation is the same on all trials.

### Optimization

To fit the model, we minimize the log-likelihood cost function

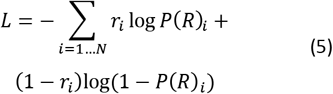

using MATLAB built-in function fmincon. At every iteration of the optimizer we evaluated equation (4), first computing values of all probability densities on a grid of 300 by 300 points in the 2d domain [*-π, π] × [-π, π*], and integrating numerically using MATLAB function *trapz* over |*x*| < |*y*| for the numerator and over the whole domain for the denominator. We ran these calculations on GPU (NVIDIA RTX 2080Ti) using MATLAB Parallel Computing Toolbox.

### Success rate with a one-sided strategy

We estimated the success rate that animals could reach when taking into account only one stimulus by first computing P(R) for every trial using a model where one concentration was set to zero and the other one to 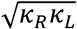 of that animal. We sampled choices using the stimulus conditions as they appeared in the experimental dataset 1000 times and computed an average percent correct over repetitions and an average across animals.

### Maximum perceptual acuity

By analogy with a 1d psychometric curve, we defined points of maximum perceptual acuity in the stimulus space as conditions (pairs of angles) where the change in *P*(*R*) was the largest for a small fixed change in the stimuli. We found these conditions from the probability surface *P*(*R*) given by the full model by computing the squared norm of the gradient vector, 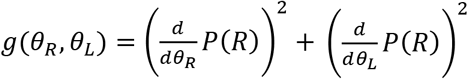 and selecting {*θ_R_,θ_L_*} conditions for which the values of *g* were in the top 5%. Among these conditions, we analyzed those with P(R) ≈ 0.5 (0.48 ≤ P(R) ≤ 0.52), which we call *maximum gradient* conditions (**Fig. 3c,h**, white) with a pooled right-choice probability of P^0.5^. For n = 28 animals this procedure gave at least 3 unique maximum gradient conditions. For n = 12 animals, the initial criterion gave fewer than three maximum gradient conditions, and we expanded the allowed range to have at least 3: we set (0.47 ≤ P(R) ≤ 0.53) for n = 7 animals, (0.46 ≤ P(R) ≤ 0.54) for n = 1, (0.42 ≤ P(R) ≤ 0.58) for n = 1, (0.40 ≤ P(R) ≤ 0.60) for n = 1, (0.38 ≤ P(R) ≤ 0.62) for n = 1, and (0.28 ≤ P(R) ≤ 0.72) for n = 1.

We then determined the neighboring conditions by changing one orientation at a time by 9°, which resulted in an increase (“+”) or decrease (“-”) of P(R) relative to P^0.5^ (**Fig. 3c**). For example, P^R-^ corresponded to the probability of right choice pooled from all conditions in which *θ_R_* changed relative to maximum gradient conditions in the direction of P(R) decrease. Here, the stimulus space was binned to a 9° grid. In a separate analysis, for an animal with 3° condition binning, we changed both orientations simultaneously by +-3°, “along” and “against” the gradient of P(R), and obtaining P^+^ and P^+^ respectively (**Fig. 3h**).

We tested that probabilities in the neighboring conditions (P^L+^, P^R+^, P^L-^, P^R-^ in case of 9°-binned conditions, and P^+^, P^-^ in case of 3°-binned conditions) were significantly different from maximum gradient probabilities P^0.5^ using a two-tailed *χ*^2^ test with df = 1, and doing pairwise comparisons of right choice frequencies, with a correction for multiple comparisons. For a population summary (**Fig. 3e**) we computed P^L+^, P^R+^, P^L-^, P^R-^ with increasing angle increments of 9°, 18°, and 27° and reported the cumulative number of animals for which at least one of the four probabilities was significantly different from P^0.5^, using a two-tailed *χ*^2^ test with df = 1 and a criterion α=0.05/4.

### History biases during high and low engagement

We first identified periods of high and low engagement in every session. For a given session, we computed a running estimate of the success rate in a sliding window of 10 trials (average performance in the window was assigned to the last trial of that window). We centered the running estimate by subtracting the mean success rate of the session. All trials with the centered success rate estimate exceeding a fixed threshold of 10% were labeled as high engagement, and all trials in which the centered success rate estimate was lower than −10% were labeled as low engagement. We confirmed the stability of our results using threshold values of 5%, 15%, and 20% (data not shown). When identifying engagement epochs, time-out trials were counted as failures, but we discarded these trials for all the analysis that followed, consistently with the rest of this study.

Next, we computed the log-likelihood *L* of outcomes in high- and low-engagement trials (*r^h^* and *r^l^* respectively) given the probabilities predicted by the full model that accounted for trial history, and by a history-free model fitted separately (*p_h_* and *p*_0_ respectively) (see Methods: Model Design). For binary outcomes *r* and model-derived probabilities *p*, we computed trial-wise the log-likelihood using the formula *L*(*r, p*) = *r* log(*p*) + (1 – *r*) log(1 – *p*) with stimulus conditions binned to a 9° grid. Applying two different trial selections and two different models we obtained *L*(*r^h^, p_h_*) for the loglikelihoods of high-engagement trial outcomes given the model with history, *L*(*r^l^,p_h_*) for the log-likelihoods of low-engagement trial outcomes given the model with history, *L*(*r^h^*, *p*_0_) for the loglikelihoods of high-engagement trials given the history-free model, and *L*(*r^l^*, *p*_0_) for the log-likelihoods of low-engagement trials given the history-free model. We next computed the differences of log-likelihood averages between models with and without history terms, using high-engagement trials, Δ*L_h_* = 〈*L*(*r^h^*, *p_h_*)〉 – 〈*L*(*r^h^*, *p*_0_)〉 and low-engagement trials, Δ*L_t_* = 〈*L*(*r^l^*, *p_h_*)〉 – 〈*L*(*r^l^*, *p*_0_)〉, (**Fig. 4d**).

Next, we computed the average of each of these log-likelihoods across all trials for every pair of orientations (*θ_L_, θ_R_*) thus obtaining maps of 〈*L*(*r*\*,*p_*_)〈_θ_* as a function of orientations (*θ_L_,θ_R_*). We discarded any stimulus conditions where the number of trials was < 10. We computed the difference between history-dependent and history-free maps of 〈*L*(*r*\*,*p*_*_)〉_*θ*_ separately for high- and low-performance trials, i.e. Δ*L_hθ_* = 〈*L*(*r^h^*, *p*_h_)〉_*θ*_ – 〈*L*(*r^h^*, *p*_h_)〉_*θ*_ and Δ*L_lθ_* = 〈*L*(*r^l^*, *p*_h_)〉_*θ*_ – 〈*L*(*r^l^*, *p*_0_)〉_*θ*_, and for all trials together, Δ*L_θ_* = 〈*L*(*y, p_h_*)〉_*θ*_ – 〈*L*(*y,p*_0_)〉_*θ*_. For a population summary (**Fig. 4e**) of Δ*L_θ_*, Δ*L_hθ_*, and Δ*L_lθ_*, we normalized Δ*L\** maps of every animal by the standard deviation across all stimulus conditions, and averaged the resulting maps across animals.

### Model comparison

#### AIC

We compared the cumulative Gaussian psychometric model to our history-free model, and the history-free model to the model with history priors, using the Akaike Information Criterion (AIC) defined as *AIC* = –2*L* + 2*k* where *k* is the number of parameters (4 for Gaussian model, 5 for the history-free model, 8 for model with history) and *L* is the log-likelihood value of the best fit. We computed *L* using the binomial log-likelihood formula

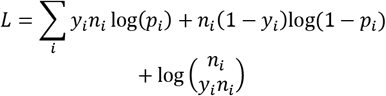

where *i* corresponds to a 9°-binned unique stimulus condition defined by (*θ_L_, θ_R_*) for the history-free to Gaussian model comparison and (*θ_L_,θ_R_,r_h_,s_h_*) for the history-free to the history-dependent model comparison, *y_i_* is the proportion of successes, *n_i_* is the total number of trials, and *p_i_* is the success rate given by either one of three models. We computed and reported Δ*AIC* = *AIC* (*Gauss*) – *AIC*(*HistFree*), and Δ*AIC* = *AIC* (*HistFree*) – *AIC*(*HistDependent*) for the final quantification.

#### Fraction of explained deviance

To estimate how much explanatory power is gained by fitting the history-free model in comparison to the Gaussian psychometric model, and by the history-dependent model in comparison to the history-free model, we computed the fraction of explained deviance. Deviance is defined as two times the log of the ratio of the saturated model likelihood *l*(*θ_max_*; *y*) to optimal model likelihood 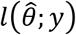

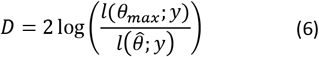

where *y* are observations, 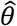 are estimated parameters, and *θ_max_* are parameters of the saturated model.

For binomial data, deviance is

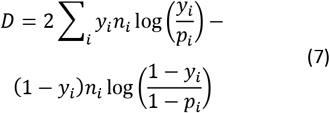

where *y_i_n_i_* is the number of successes for stimulus condition *i*, is the number of trials, and *p_i_* is the probability of success in condition *i* given by the fitted model with parameters 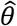. For the cumulative Gaussian psychometric function *ψ*(Δ*θ*; *α,β,γ,λ*) a stimulus condition is defined by a pair of angles {*θ_R_, θ_L_*} in a history-free model, and a pair of angles with trial history {*θ_R_,θ_L_,s_h_,r_h_*} in a model with history.

We first computed the deviance of the null model, with the same P(R) = *p_null_* rate for all conditions (computed as a grand average P(R) across trials). We then used the formula for deviance *D* (7), with *p_i_* = *P_null_* when computing null deviance *D_null_*, *p_i_* = *p_i_*(*HF*) as predicted by history-free model when computing history-free deviance *D_HF_*, *p_i_* = *p_i_*(*HD*) as predicted by the history-dependent model when computing history-dependent deviance *D_HD_*, and *p_i_* = *p_i_*(*Gauss*) as predicted by the Gaussian model when computing Gaussian deviance *D_Gauss_*. Here a condition *i* corresponded to a unique pair of orientations (*θ_L_, θ_R_*) when comparing the Gaussian model with the history-free model, and to a pair of orientations together with history inputs (*θ_L_, θ_R_, s_h_, r_h_*) when comparing the history-free model and the history-dependent model; the fraction of right choices *y_i_* and the total number of trials per condition *n_i_* changed accordingly. We computed the fraction of explained deviance (*FDE*) for the three models as *FDE_HF_* = 100% · (*D_null_* – *D_HF_*)/*D_null_*, *FDE_HD_* = 100% · (*D_null_* – *D_HD_*)/*D_null_* and *FDE_Gauss_* = 100% · (*D_null_* – *D_Gauss_*)/ *D_null_*, and finally we computed difference in the fraction of deviance explained as Δ*FDE* = *FDE_HF_* – *FDE_Gauss_* or Δ*FDE* = *FDE_HD_* – *FDE_HF_*. For this analysis, we trained each model on 50% randomly sampled trials and computed deviances from the other 50% of trials. We tested the significance of Δ*FDE* > 0 for a population of animals using the Wilcoxon signed rank test.

---

**Supplementary Figure 1.**
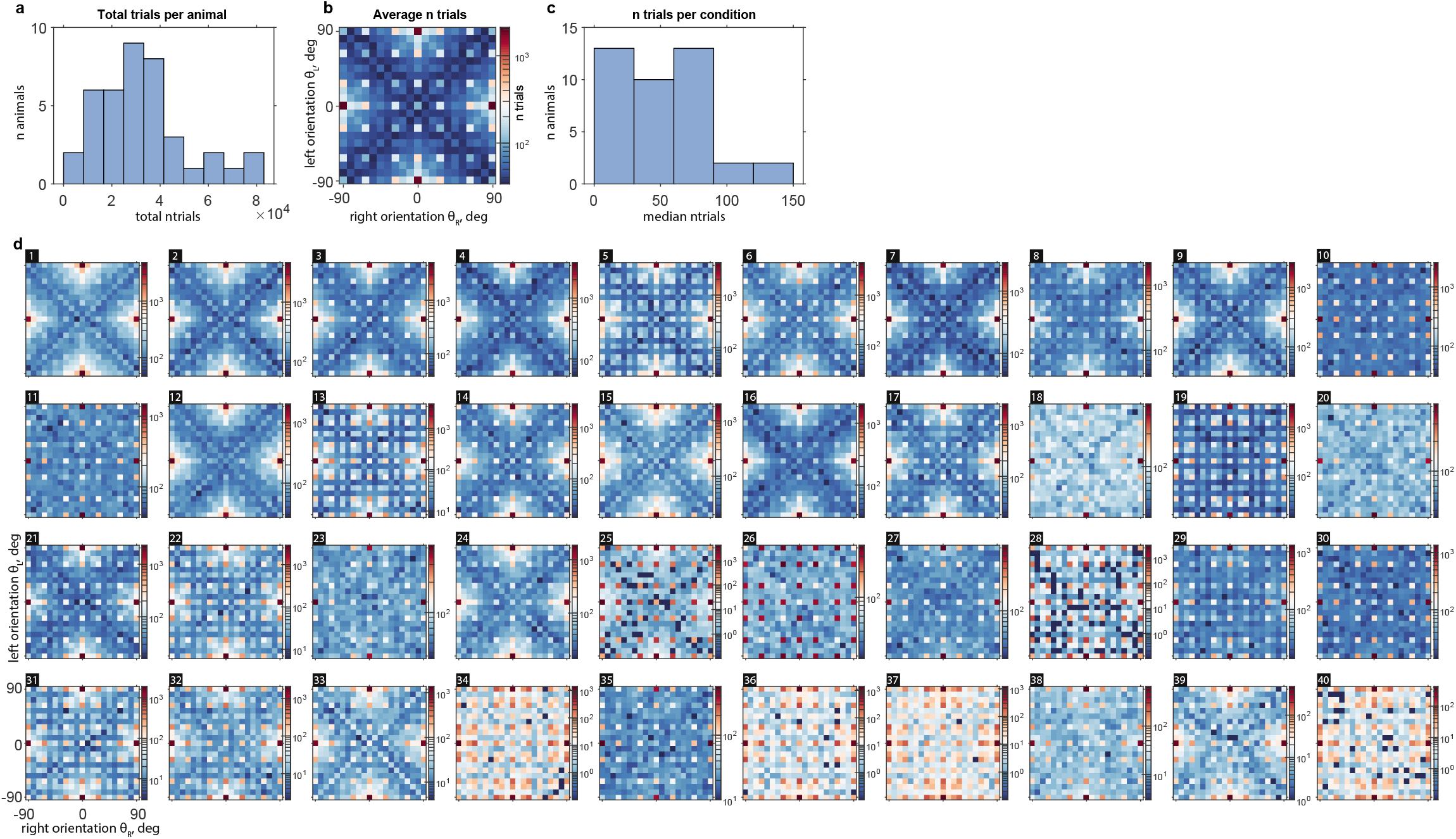
A high number of trials was collected from n = 40 animals. **a.** Total number of trials collected for all animals. **b.** Population-average number of trials for every stimulus condition (pair of angles); color bar – number of trials, log scale. **c.** Median number of trials across conditions for every animal. **d.** Number of trials for every stimulus condition and every animal, axes as in b; number in black square – animal ID, the same as in Supplementary Figure 3 and Supplementary Table 1.

**Supplementary Figure 2.**
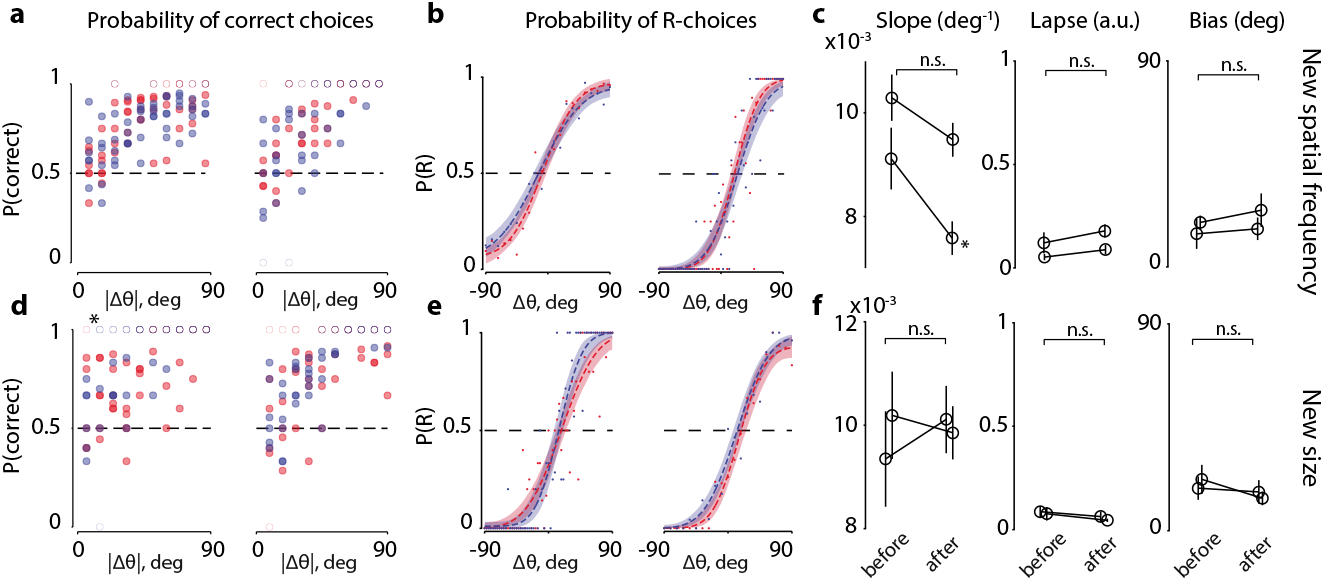
Performance is invariant relative to size and spatial frequency transformations of the stimulus. **a.** Correct rate as a function of trial easiness |Δθ|, comparing 3 sessions (divided into 6 groups of trials—one dot per group, dots may overlap—with ~75 trials/group) before a change in spatial frequency of the gratings (red dots) and after the change (blue dots) (spatial frequency, SF = 0.008 −> 0.016 cpd for mouse A, left panel, and SF = 0.0016 −> 0.032 cpd for mouse B, right panel). Open circles for correct rates {0, 1}. Data for the right panel (mouse B, minimal angular difference 3°) was grouped into 9° bins to improve visualization. For statistical comparison, we compared binned data (non-overlapping 18° bins) from before vs after conditions and found no significant difference (p > 0.05, Wilcoxon rank-sum test). **b.** Psychometric curves from 3 sessions before (red) and after (blue) changing the spatial frequency of stimuli. Same data as in a, used as right/left choices; dots for average P(R) as in the data; dotted lines for the fits; colored bands for bootstrap confidence intervals. **c.** Comparison of fitting parameters: slope, lapse rate, and bias, before and after changing spatial frequency of stimuli (mean ± s.e.m., n = 2 mice, n.s. for p > 0.05, and ‘*’ for p < 0.05, unpaired t-test for individual animals, paired t-test for comparison across animals). **d-f.** Same as a-c, but for changes in stimulus size (20° −> 25° visual angle, n = 2 mice B, C). Data sampled at 3° angle difference has been grouped into 9° bins to improve visualization. Left panels: statistical difference for |△θ| bin = 18° (‘*’ for p < 0.05) reflects an improvement in the performance after changing stimulus size.

**Supplementary Figure 3.**
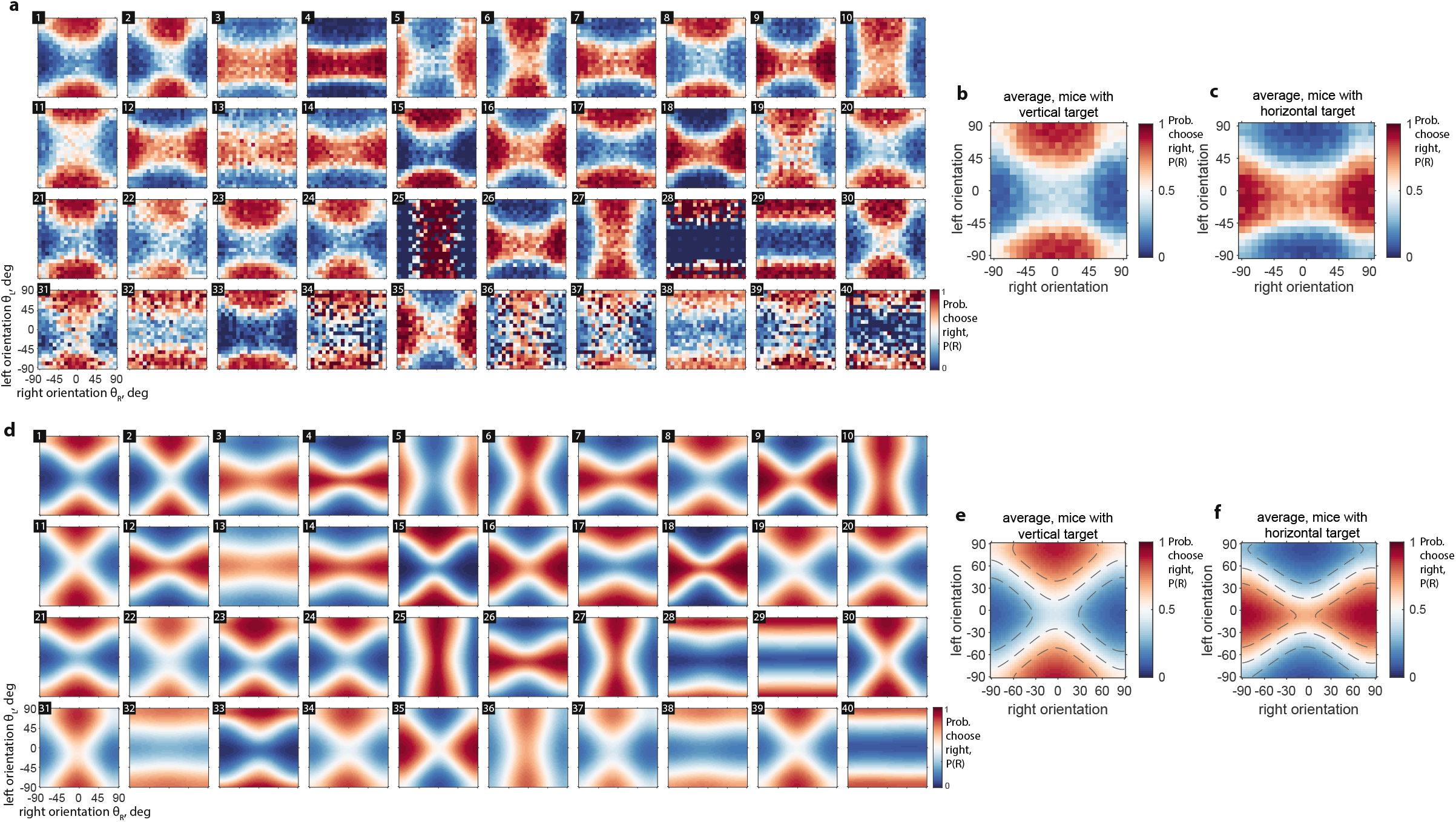
Choices of mice are largely determined by the rewarded side in the two-dimensional stimulus space, and choice model recapitulates choice probability. **a.** Probability of right choice, P(R), for all mice. Stimulus conditions are binned to 9°. Color limits are the same in all panels and in b-c. Animal IDs (number in a black square) are as in Supplementary Figure 1 and Supplementary Table 1. **b.** Average P(R) across animals trained to find a more vertical orientation. **c.** Average P(R) across animals trained to find a more horizontal orientation. **d.** Model P(R) surfaces for every animal, same color bar on all panels, and as in e and f. **e.** Average P(R) surface of all animals trained to find the more vertical target. Dashed lines at P(R) values of 0.25, 0.5, 0.75. **f.** Average P(R) surface of all animals trained to find the more horizontal target.

**Supplementary Figure 4.**
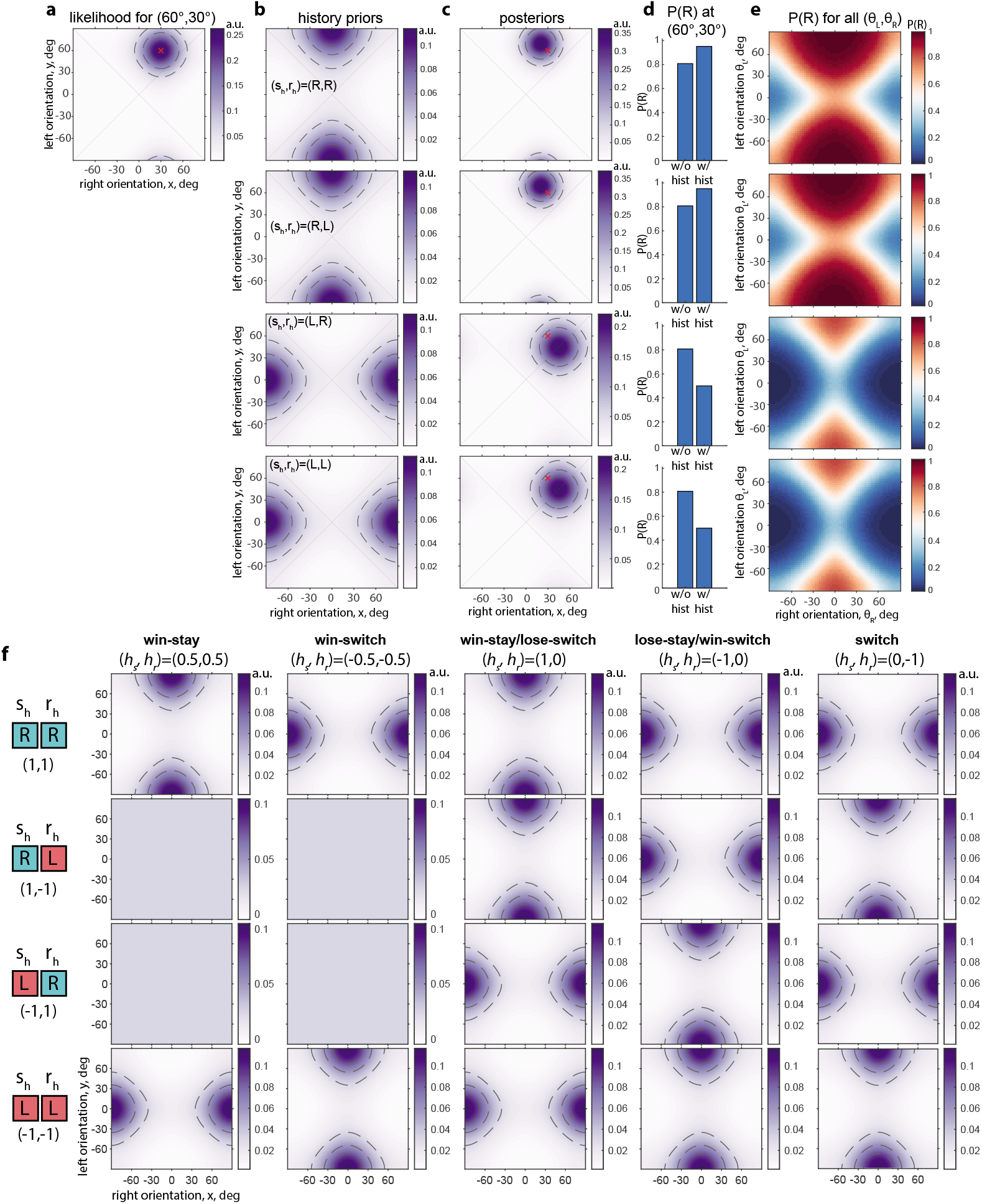
History priors can represent many possible strategies, and affect choice probability, P(R), by shifting the probability density p(x, y) inside or outside the |x| < |y| region **a.** Probability density (p.d., shown by color saturation) p(x, y) (Methods, Eq. 1) induced by stimuli (θ_R_, θ_L_) = (30°, 60°) (red cross) in a model with κ_R_ = κ_L_ = 2, b_R_ = b_L_ = 0, and κ_b_ = 0; dashed lines show distribution quartiles. **b.** History prior p_h_(x, y) (Methods, Eq. 3) corresponding to the win-stay/lose-switch strategy, (h_s_, h_r_) = (0, 1), with κ_h_ = 5, and four possible target-response combinations (s_h_, r_h_) on the previous trial. Top to bottom: (s_h_, r_h_) = (R, R); (R, L); (L, R); (L, L). **c.** Posterior p.d. p(θR*, θ_L_*): normalized product of p(x, y) and p_h_(x, y) before integration over |x| – |y|, with (s_h_, r_h_) same as in b in the same row. **d.** probability of right choice P(R) for (θ_R_, θ_L_)=(30°, 60°) with and without history bias. **e.** P(R) with strategy for all (θ_R_, θ_L_) corresponding to (s_h_, r_h_) in b in the same row. **f.** History priors p_h_(x, y) for five example history-based strategies (columns) shown for all four possible combinations of target and choice on the previous trial (rows); κ_h_ = 1 in all cases.

**Supplementary Figure 5.**
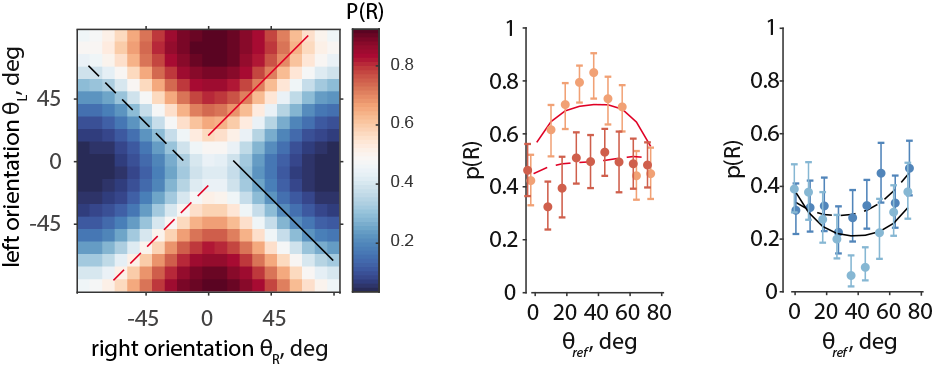
Variation of P(R) with reference orientation θ_ref_ is larger in the data than in the model. Left. Example mouse, selected here for its low translational bias and approximately equal concentrations for right and left stimuli, which results in a regularly shaped P(R) dependency on θ_ref_ (cf. Figure 2c). Center and right. P(R) along θ_ref_ for the △θ = const conditions marked on the left panel, as predicted by the model (lines) and as in the data (dots with whiskers). Dots of a lighter shade (orange, light blue) correspond to the solid lines.

**Supplementary Table 1.**
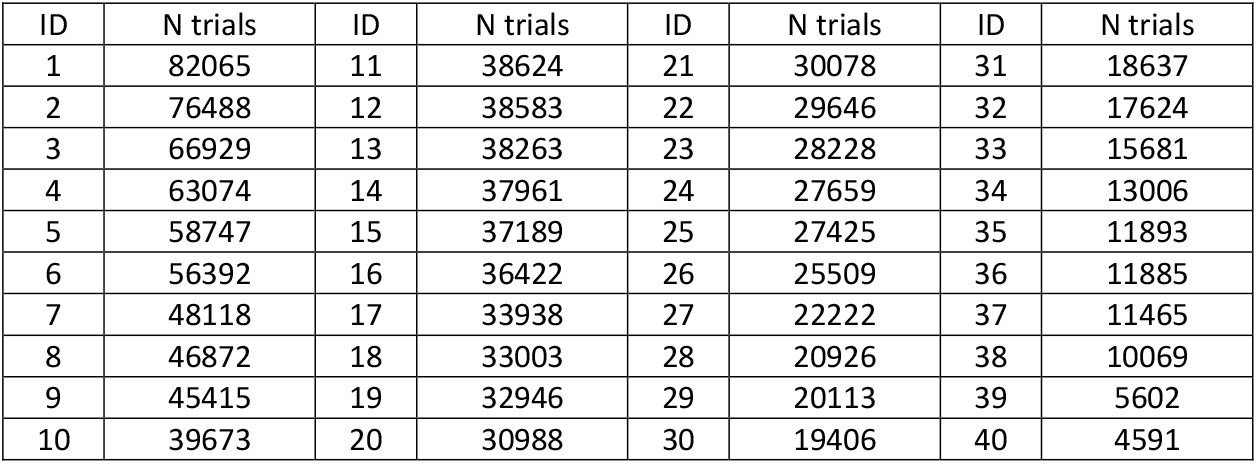
Total number of trials per animal. “ID” columns show animal identification numbers as in Supplementary Figures 1 and 3, “N Trials” columns show total number of trials of the corresponding mouse used in the analysis throughout the paper.

| ID | N trials | ID | N trials | ID | N trials | ID | N trials |
| --- | --- | --- | --- | --- | --- | --- | --- |
| 1 | 82065 | 11 | 38624 | 21 | 30078 | 31 | 18637 |
| 2 | 76488 | 12 | 38583 | 22 | 29646 | 32 | 17624 |
| 3 | 66929 | 13 | 38263 | 23 | 28228 | 33 | 15681 |
| 4 | 63074 | 14 | 37961 | 24 | 27659 | 34 | 13006 |
| 5 | 58747 | 15 | 37189 | 25 | 27425 | 35 | 11893 |
| 6 | 56392 | 16 | 36422 | 26 | 25509 | 36 | 11885 |
| 7 | 48118 | 17 | 33938 | 27 | 22222 | 37 | 11465 |
| 8 | 46872 | 18 | 33003 | 28 | 20926 | 38 | 10069 |
| 9 | 45415 | 19 | 32946 | 29 | 20113 | 39 | 5602 |
| 10 | 39673 | 20 | 30988 | 30 | 19406 | 40 | 4591 |

## References

Abbott, L.F., Bock, D.D., Callaway, E.M., Denk, W., Dulac, C., Fairhall, A.L., Fiete, I., Harris, K.M., Helmstaedter, M., Jain, V., et al. (2020). The Mind of a Mouse. Cell 182, 1372–1376.

Abdolrahmani, M., Lyamzin, D.R., Aoki, R., and Benucci, A. (2021). Attention Decorrelates Sensory and Motor Signals in the Mouse Visual Cortex. BioRxiv 615229.

Abrahamyan, A., Silva, L.L., Dakin, S.C., Carandini, M., and Gardner, J.L. (2016). Adaptable history biases in human perceptual decisions. Proc. Natl. Acad. Sci. U. S. A. 113, E3548–57.

Akrami, A., Kopec, C.D., Diamond, M.E., and Brody, C.D. (2018). Posterior parietal cortex represents sensory history and mediates its effects on behaviour. Nature 554, 368–372.

Alemi-Neissi, A., Rosselli, F.B., and Zoccolan, D. (2013). Multifeatural Shape Processing in Rats Engaged in Invariant Visual Object Recognition. J. Neurosci. 33, 5939 LP – 5956.

Andermann, M., Kerlin, A., and Reid, C. (2010). Chronic cellular imaging of mouse visual cortex during operant behavior and passive viewing. Front. Cell. Neurosci. 4, 3.

Aoki, R., Tsubota, T., Goya, Y., and Benucci, A. (2017). An automated platform for high-throughput mouse behavior and physiology with voluntary head-fixation. Nat. Commun. 8, 1196.

Ashwood, Z.C., Roy, N.A., Stone, I.R., Churchland, A.K., Pouget, A., and Pillow, J.W. (2020). Mice alternate between discrete strategies during perceptual decision-making. BioRxiv 2020.10.19.346353.

Beck, J.M., Ma, W.J., Pitkow, X., Latham, P.E., and Pouget, A. (2012). Not noisy, just wrong: the role of suboptimal inference in behavioral variability. Neuron 74, 30–39.

Britten, K.H., Newsome, W.T., Shadlen, M.N., Celebrini, S., and Movshon, J.A. (1996). A relationship between behavioral choice and the visual responses of neurons in macaque MT. Vis. Neurosci. 13, 87–100.

Brunton, B.W., Botvinick, M.M., and Brody, C.D. (2013). Rats and Humans Can Optimally Accumulate Evidence for Decision-Making. Science (80-.). 340, 95 LP – 98.

Burgess, C.P., Lak, A., Steinmetz, N.A., Zatka-Haas, P., Bai Reddy, C., Jacobs, E.A.K., Linden, J.F., Paton, J.J., Ranson, A., Schröder, S., et al. (2017). High-Yield Methods for Accurate Two-Alternative Visual Psychophysics in Head-Fixed Mice. Cell Rep. 20, 2513–2524.

Busse, L., Ayaz, A., Dhruv, N.T., Katzner, S., Saleem, A.B., Schölvinck, M.L., Zaharia, A.D., and Carandini, M. (2011). The detection of visual contrast in the behaving mouse. J. Neurosci. 31, 11351–11361.

Constantinople, C.M., Piet, A.T., and Brody, C.D. (2019). An Analysis of Decision under Risk in Rats. Curr. Biol. 29, 2066–2074.e5.

Corrado, G.S., Sugrue, L.P., Seung, H.S., and Newsome, W.T. (2005). Linear-Nonlinear-Poisson models of primate choice dynamics. J. Exp. Anal. Behav. 84, 581–617.

Dakin, S.C., Tibber, M.S., Greenwood, J.A., Kingdom, F.A.A., and Morgan, M.J. (2011). A common visual metric for approximate number and density. Proc. Natl. Acad. Sci. 108, 19552 LP – 19557.

DiCarlo, J.J., and Cox, D.D. (2007). Untangling invariant object recognition. Trends Cogn. Sci. 11, 333–341.

DiCarlo, J.J., Zoccolan, D., and Rust, N.C. (2012). How does the brain solve visual object recognition? Neuron 73, 415–434.

Dosher, B.A., and Lu, Z.L. (1998). Perceptual learning reflects external noise filtering and internal noise reduction through channel reweighting. Proc. Natl. Acad. Sci. U. S. A. 95, 13988–13993.

Drugowitsch, J., Wyart, V., Devauchelle, A.-D., and Koechlin, E. (2016). Computational Precision of Mental Inference as Critical Source of Human Choice Suboptimality. Neuron 92, 1398–1411.

Fründ, I., Wichmann, F.A., and Macke, J.H. (2014). Quantifying the effect of intertrial dependence on perceptual decisions. J. Vis. 14.

Gardner, J.L. (2019). Optimality and heuristics in perceptual neuroscience. Nat. Neurosci. 22, 514–523.

Girshick, A.R., Landy, M.S., and Simoncelli, E.P. (2011). Cardinal rules: visual orientation perception reflects knowledge of environmental statistics. Nat. Neurosci. 14, 926–932.

Glickfeld, L.L., Histed, M.H., and Maunsell, J.H.R. (2013). Mouse Primary Visual Cortex Is Used to Detect Both Orientation and Contrast Changes. 33, 19416–19422.

Goard, M.J., Pho, G.N., Woodson, J., and Sur, M. (2016). Distinct roles of visual, parietal, and frontal motor cortices in memory-guided sensorimotor decisions. Elife 5, 1–30.

Hernández, A., Salinas, E., García, R., and Romo, R. (1997). Discrimination in the Sense of Flutter: New Psychophysical Measurements in Monkeys. J. Neurosci. 17, 6391 LP – 6400.

Hubel, D.H., and Wiesel, T.N. (1962). Receptive fields, binocular interaction and functional architecture in the cat’s visual cortex. J. Physiol. 160, 106–154.

Huberman, A.D., and Niell, C.M. (2011). What can mice tell us about how vision works? Trends Neurosci. 34, 464–473.

Jazayeri, M., and Movshon, J.A. (2007). A new perceptual illusion reveals mechanisms of sensory decoding. Nature 446, 912–915.

Jin, M., Beck, J.M., and Glickfeld, L.L. (2019). Neuronal Adaptation Reveals a Suboptimal Decoding of Orientation Tuned Populations in the Mouse Visual Cortex. J. Neurosci. 39, 3867 LP – 3881.

Jogan, M., and Stocker, A.A. (2014). A new two-alternative forced choice method for the unbiased characterization of perceptual bias and discriminability. J. Vis. 14, 20.

Jurjut, O., Georgieva, P., Busse, L., and Katzner, S. (2017). Learning Enhances Sensory Processing in Mouse V1 before Improving Behavior. J. Neurosci. 37, 6460–6474.

Krechevsky, I. (1938). An experimental investigation of the principle of proximity in the visual perception of the rat. J. Exp. Psychol. 22, 497–523.

Lak, A., Hueske, E., Hirokawa, J., Masset, P., Ott, T., Urai, A.E., Donner, T.H., Carandini, M., Tonegawa, S., Uchida, N., et al. (2020). Reinforcement biases subsequent perceptual decisions when confidence is low, a widespread behavioral phenomenon. Elife 9, e49834.

Laquitaine, S., and Gardner, J.L. (2018). A Switching Observer for Human Perceptual Estimation. Neuron 97, 462–474.e6.

Lashley, K.S. (1938). The Mechanism of Vision: XV. Preliminary Studies of the Rat’s Capacity for Detail Vision. J. Gen. Psychol. 18, 123–193.

Long, M., Jiang, W., Liu, D., and Yao, H. (2015). Contrast-dependent orientation discrimination in the mouse. Sci. Rep. 5, 15830.

Luo, L., Callaway, E.M., and Svoboda, K. (2018). Genetic Dissection of Neural Circuits: A Decade of Progress. Neuron 98, 256–281.

Madisen, L., Garner, A.R., Shimaoka, D., Chuong, A.S., Klapoetke, N.C., Li, L., van der Bourg, A., Niino, Y., Egolf, L., Monetti, C., et al. (2015). Transgenic Mice for Intersectional Targeting of Neural Sensors and Effectors with High Specificity and Performance. Neuron 85, 942–958.

Martinho, A. 3rd, and Kacelnik, A. (2016). Ducklings imprint on the relational concept of “same or different”. Science 353, 286–288.

McGinley, M.J., Vinck, M., Reimer, J., Batista-Brito, R., Zagha, E., Cadwell, C.R., Tolias, A.S., Cardin, J.A., and McCormick, D.A. (2015). Waking State: Rapid Variations Modulate Neural and Behavioral Responses. Neuron 87, 1143–1161.

Odoemene, O., Pisupati, S., Nguyen, H., and Churchland, A.K. (2018). Visual Evidence Accumulation Guides Decision-Making in Unrestrained Mice. J. Neurosci. 38, 10143 LP – 10155.

Pho, G.N., Goard, M.J., Woodson, J., Crawford, B., and Sur, M. (2018). Task-dependent representations of stimulus and choice in mouse parietal cortex. Nat. Commun. 9, 2596.

Pinto, L., Goard, M.J., Estandian, D., Xu, M., Kwan, A.C., Lee, S.-H., Harrison, T.C., Feng, G., and Dan, Y. (2013). Fast modulation of visual perception by basal forebrain cholinergic neurons. Nat. Neurosci. 16, 1857–1863.

Pinto, L., Koay, S.A., Engelhard, B., Yoon, A.M., Deverett, B., Thiberge, S.Y., Witten, I.B., Tank, D.W., and Brody, C.D. (2018). An Accumulation-of-Evidence Task Using Visual Pulses for Mice Navigating in Virtual Reality. Front. Behav. Neurosci. 12, 1–19.

Pinto, L., Rajan, K., DePasquale, B., Thiberge, S.Y., Tank, D.W., and Brody, C.D. (2019). Task-Dependent Changes in the Large-Scale Dynamics and Necessity of Cortical Regions. Neuron 104, 810–824.e9.

Poort, J., Khan, A.G., Pachitariu, M., Nemri, A., Orsolic, I., Krupic, J., Bauza, M., Sahani, M., Keller, G.B., Mrsic-Flogel, T.D., et al. (2015). Learning Enhances Sensory and Multiple Non-sensory Representations in Primary Visual Cortex. Neuron 86, 1478–1490.

Prusky, G.T., and Douglas, R.M. (2004). Characterization of mouse cortical spatial vision. Vision Res. 44, 3411–3418.

Reimer, J., McGinley, M.J., Liu, Y., Rodenkirch, C., Wang, Q., McCormick, D.A., and Tolias, A.S. (2016). Pupil fluctuations track rapid changes in adrenergic and cholinergic activity in cortex. Nat. Commun. 7, 13289.

Renart, A., and Machens, C.K. (2014). Variability in neural activity and behavior. Curr. Opin. Neurobiol. 25, 211–220.

Resulaj, A., Ruediger, S., Olsen, S.R., and Scanziani, M. (2018). First spikes in visual cortex enable perceptual discrimination. Elife 7, e34044.

Reuter, J.H. (1987). Tilt discrimination in the mouse. Behav. Brain Res. 24, 81–84.

Riesenhuber, M., and Poggio, T. (2000). Models of object recognition. Nat. Neurosci. 3, 1199–1204.

Romo, R., Brody, C.D., Hernández, a, and Lemus, L. (1999). Neuronal correlates of parametric working memory in the prefrontal cortex. Nature 399, 470–473.

Runyan, C.A., Piasini, E., Panzeri, S., and Harvey, C.D. (2017). Distinct timescales of population coding across cortex. Nature 548, 92–96.

Scott, B.B., Constantinople, C.M., Erlich, J.C., Tank, D.W., and Brody, C.D. (2015). Sources of noise during accumulation of evidence in unrestrained and voluntarily head-restrained rats. Elife 4, e11308.

Steinmetz, N.A., Zatka-haas, P., Carandini, M., and Harris, K.D. (2019). Distributed coding of choice, action and engagement across the mouse brain. Nature.

Stocker, A.A., and Simoncelli, E.P. (2006). Noise characteristics and prior expectations in human visual speed perception. Nat. Neurosci. 9, 578–585.

Urai, A.E., Braun, A., and Donner, T.H. (2017). Pupil-linked arousal is driven by decision uncertainty and alters serial choice bias. Nat. Commun. 8, 14637.

Vermaercke, B., and Op de Beeck, H.P. (2012). A Multivariate Approach Reveals the Behavioral Templates Underlying Visual Discrimination in Rats. Curr. Biol. 22, 50–55.

Wang, L., Rangarajan, K. V, Gerfen, C.R., and Krauzlis, R.J. (2018). Activation of Striatal Neurons Causes a Perceptual Decision Bias during Visual Change Detection in Mice. Neuron 97, 1369–1381.e5.

Wang, L., McAlonan, K., Goldstein, S., Gerfen, C.R., and Krauzlis, R.J. (2020). A Causal Role for Mouse Superior Colliculus in Visual Perceptual Decision-Making. J. Neurosci. 40, 3768 LP – 3782.

Whiteley, L., and Sahani, M. (2012). Attention in a Bayesian Framework. Front. Hum. Neurosci. 6, 100.

Wichmann, F.A., and Hill, N.J. (2001). The psychometric function: I. Fitting, sampling, and goodness of fit. Percept. Psychophys. 63, 1293–1313.

Wyart, V., and Koechlin, E. (2016). Choice variability and suboptimality in uncertain environments. Curr. Opin. Behav. Sci. 11, 109–115.

You, W.-K., and Mysore, S.P. (2020). Endogenous and exogenous control of visuospatial selective attention in freely behaving mice. Nat. Commun. 11, 1986.

Yu, A.J., and Cohen, J.D. (2008). Sequential effects: Superstition or rational behavior? In Advances in Neural Information Processing Systems, D. Koller, D. Schuurmans, Y. Bengio, and L. Bottou, eds. (Curran Associates, Inc.), pp. 1873–1880.

Zoccolan, D., Cox, D.D., and Benucci, A. (2015). Editorial: What can simple brains teach us about how vision works. Front. Neural Circuits 9, 51.

